# Active density fluctuations in bacterial binary mixtures

**DOI:** 10.1101/2023.05.10.540167

**Authors:** Silvia Espada Burriel, Remy Colin

## Abstract

In wild environments, physical and biochemical interactions between intermixed motile and sessile microorganisms give rise to spatial organization that is key for the functioning and ecology of complex communities. However, how motility-driven physical interactions contribute to shaping multispecies communities remains little understood. To address this gap, we investigated model binary mixtures of motile and non-motile *Escherichia coli* bacteria. We discovered a new type of non-equilibrium self-organization, wherein large-scale density fluctuations of non-motile bacteria emerge when mixed with motile ones under physiologically relevant conditions. Systematically exploring the phase diagram in microfluidics experiments and combining them with modeling and simulations, we uncovered the two-pronged physical mechanism of emergence: Circular swimming of motile cells close to surfaces generates recirculating hydrodynamic flows that advect non-motile cells, while sedimentation, by breaking the vertical symmetry, is essential for their local accumulation. This active self-organization behavior in mixed bacterial populations appears crucial for complex microbial community structuration.

## Introduction

Natural microorganism communities are composed of many different species with diverse phenotypes, and they are observed to organize in a wide variety of complex spatial structures ^1–3^. This complex organization is thought to be a prominent determinant for the coexistence, ecology and evolution of multi-species communities ^2–4^. Biochemical interactions between species are known to shape the structure of microbial communities, especially when the colony forms a sessile (non-motile) surface attached biofilm ^5–8^. While having been much less investigated, physical interactions between phenotypically different species might also matter strongly for population organization, particularly where active motility is involved.

Many bacterial species, such as the model organism *Escherichia coli*, are indeed motile. They rotate helical flagella to propel themselves at speeds of∼10−100 *µ*m/s. At the single cell level, the bacterium alternates seconds-long periods of straight runs (modulo rotational Brownian motion) with short reorientations, the tumbles, to describe a random walk that allows each cell to explore the environment^9^. This motility is coupled to a sensory system, the chemotaxis pathway, which modulates the tumbling frequency according to perceived changes in physicochemical environmental conditions (nutrients, pH, temperature, etc.) to bias the cell motion towards “preferred” conditions and thus navigate the environment ^10–14^.

The constant mechanical energy input due to swimming drives bacterial suspensions out of equilibrium. In the resulting active fluids, a host of self-organization behaviors may take place, from the single cell to population levels ^15^. The bacteria can be thought of as force dipoles in a low Reynolds number fluid ^16^ which interact hydrodynamically with each other and with surfaces at long range via the fluid they displace, on top of steric interactions upon direct collisions ^17^. At single cell level, swimming bacteria tend to accumulate near surfaces due to both types of interactions ^18–22^. The proximity to the surface also alters their swimming behavior, making them swim in circular trajectories ^23–25^.

In homogeneous motile bacterial populations, hydrodynamic and steric interactions between the rod-shaped bacteria induce polar ordering which explain the emergence of swirling collective motion known as bacterial turbulence at high cell density (volume fraction *ϕ >* 1 %) ^26–31^. In contrast, if the cell speed decreases when density gets high, a motility induced phase separation (MIPS) can take place, where the active fluid phaseseparate in slow/dense and fast/sparse regions ^32–34^. This can arise because of steric interactions in absence of polar alignment in the case of active Brownian particles ^33,35^, but also in (polar) bacteria from density-induced biological downregulation of motility ^36^. Moreover, motility combined with chemotaxis towards quorum-sensing chemicals released by other cells can produce density patterns ^37^, ranging in bacteria from autoaggregation on a µ100 m scale ^38^ to both static and dynamic density structures on a millimeter to centimeter scale ^39–42^.

In heterogeneous mixtures of motile cells and non-motile (passive) agents, a few non-equilibrium effects have been observed. Motile cells are known to enhance the diffusion of passive particles ^43–45^, notably affecting their sedimentation ^46^. Large passive Brownian colloids may also experience effective attractive interactions when in suspension with motile cells ^47–49^. Moreover, numerical simulations of mixtures of active and passive Brownian particles have shown the emergence of spatial co-organization patterns in two dimensions, both for disks ^50–55^ and rods ^56^. The passive particles form dynamic clusters driven by the activity of the motile particles, which co-segregate, in a mechanism reminiscent of MIPS ^57^. Active particles with different mobilities also seem to demix ^58,59^. When passive particles are coupled to an active nematic, which commonly models dense active actin filament suspensions rather than bacteria ^60^, a fluctuation dominated phase ordering (FDPO) has been observed to emerge in simulations, where density fluctuations of the passive particles emerge as they surf on the two-dimensional active nematic ^61–63^. Experimentally, few studies have investigated physical self-organization in bacterial mixtures, although it has been observed when motility is coupled to biochemical interactions, e.g. when two strains cross-regulate each other’s motility ^64^ or compete for resources ^65^, when a non-motile strain recruit a motile one in its biofilm via chemotaxis ^66^ or when a non-motile strain hitchhike with a motile one ^67^. However, the purely physical effects of motility in such bacterial mixtures had hitherto not been studied.

In this paper, we investigated the physical effect of motility on the spatial organization of complex communities, using binary mixtures of bacteria, where one component is motile and the other not, as a minimal model system. We observe that large fluctuations of the non-motile cell density emerge, under the action of the motile cells, while the motile cell distribution remained homogeneous. Systematically varying the density of both motile and non-motile cells, we found that the density fluctuations happened in a wide, biologically relevant range. We tuned the swimming pattern of the motile cells by genetic modifications and found that enhanced surface localization and highly spatiotemporally organized swimming patterns at the surface enhance the density fluctuations. We further found that non-motile cell sedimentation is an essential component of the emergence of the fluctuations. The density patterns are dynamic, forming and dissolving on the time scale of minutes. We characterized the collective dynamics of the non-motile cells, which highlighted the importance of vertical recirculation and of hydrodynamic interactions between cells in the emergence of the density patterns. Our observations can be recapitulated in a mechanism that is supported by numerical simulations: Surface-localized swimmers generate three-dimensional recirculation flows leading to convective fluxes of non-motile cells. The latter fluxes combine with sedimentation, which breaks the top-down symmetry of the system and thus allows for local cell accumulation, hence giving rise to the density fluctuations.

## Results

### Emergence of large density fluctuations in bacterial binary mixtures

We study mixtures of motile and non-motile *E. coli*, which are fluorescently marked with respectively mCherry and mNeonGreen, in a microfabricated chamber of controlled height, *h* = 50 *µ*m (Methods), systematically varying the cell densities of both components of the binary mixture. All strains are stripped of their main surface adhesin Antigen 43 to prevent cell aggregation ^38^. To keep biochemical interactions as trivial as possible, the strains are genetically identical, except for the non-motile strain being a knockout of the gene *fliC*, which encodes for the flagellar filament subunit. A passive system composed of two non-motile strains tagged with mCherry and mNeonGreen is used as a control.

In some binary mixtures, we observed large spatial fluctuations of the density of the non-motile cells in presence of motile cells (Fig. 1a), whereas the corresponding passive system remains homogeneous (Fig. 1b). These large density fluctuations are characterized by the existence at steady state of dynamic patches of abnormally high and low densities of non-motile cells, which reorganize in a matter of minutes (Supplementary Movie 1). We quantified these density fluctuations using fluorescence intensity as a proxy for cell density (Methods). The spatial structure of the density of non-motile cells shows characteristics of giant fluctuations (Supplementary Figs. S1b and S1d). Correspondingly, the spatial autocorrelation of the non-motile fluorescence intensity shows a non-exponential decay at large length scales, (10−50) *µ*m, well above the 2∼*µ*m cell length (Supplementary Fig. S1e), showing long-ranged spatial correlations in these strong fluctuations of density.

**Figure 1:**
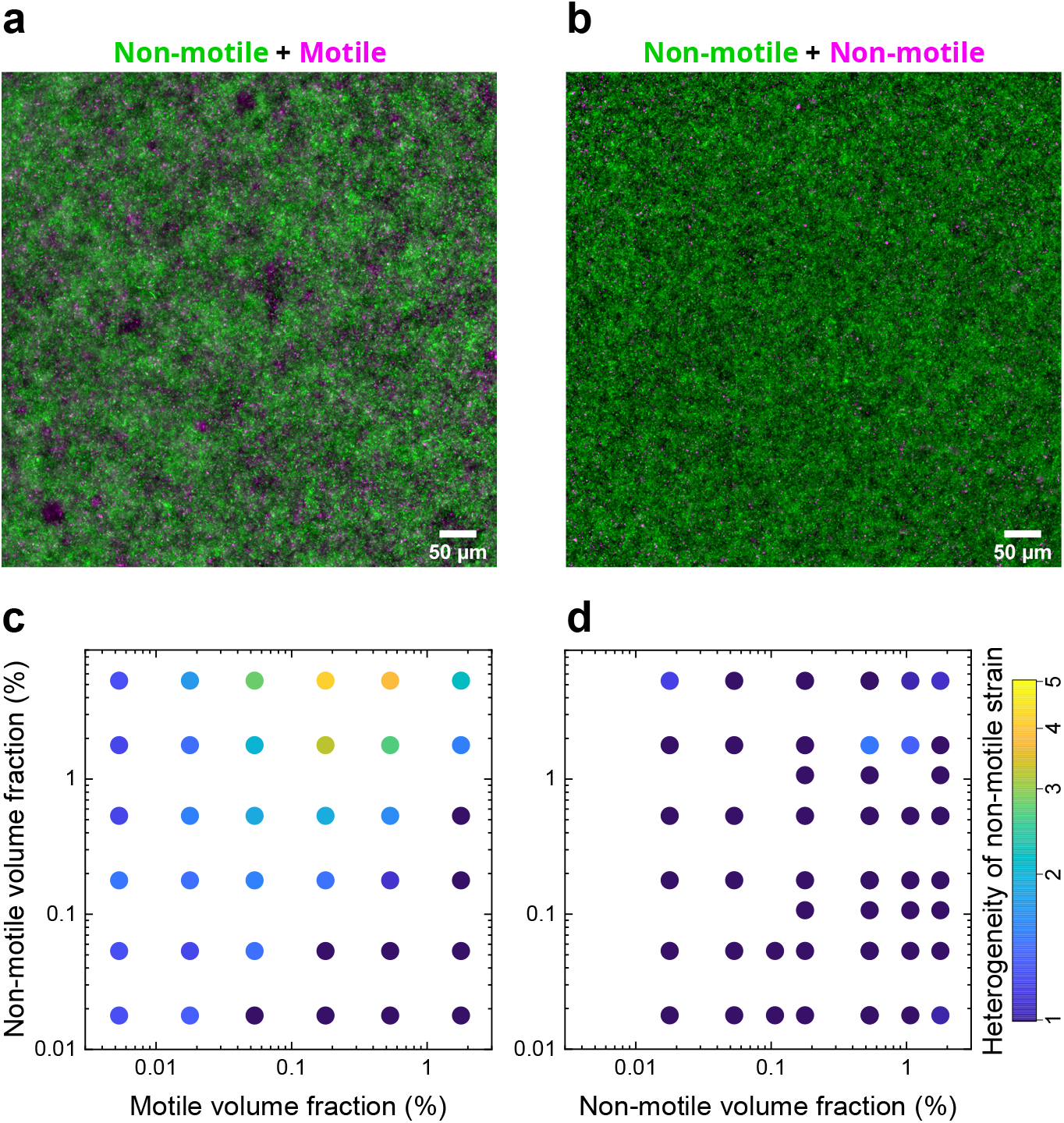
Density fluctuations in bacterial binary mixtures. a) Microscopy images of bacterial mixtures of motile strain (mCherry) and non-motile strain (mNeonGreen) of *E*.*coli* (motile cells volume fraction *ϕ*_*M*_ = 0.18%, non-motile *ϕ*_*NM*_ = 5.3%). The density of the non-motile strain exhibits strong spatial fluctuations in presence of the motile strain. b) The corresponding control where two non-motile strains of *E*.*coli* were mixed at the same volume fractions. c-d) Phase diagrams of the density fluctuations of the non-motile strain in binary mixtures of bacterial populations depending on the volume fraction of both strains. The density fluctuations are quantified as described in the Methods section by a heterogeneity level, which is represented by a color scale. Each point represents the geometric mean of at least three biological replicates. c) Mixture of a motile strain (abscissa) with a non-motile strain (ordinate). d) Mixture of two non-motile strains as a passive control system.

We used the normalized spatial variance of the cell mass on this typical length scale, called the heterogeneity level further on, as a quantification of the amplitude of the density fluctuations (Methods). During the 30 minutes after a freshly stirred mixture is introduced in the observation chamber (*t* = 0), the heterogeneity level increases until it reaches a steady state, this dynamics being accelerated when more motile cells are present (Supplementary Movie 2 and Supplementary Fig. S2). We constructed a full phase diagram of the amplitude of the fluctuations at steady state (i.e. for *t* ≃ 1 hour) as a function of the volume fractions of motile (*ϕ*_*M*_) and non-motile (*ϕ*_*NM*_) cells (Fig. 1c). The density fluctuations are observed only when non-motile cells are in a majority (*ϕ*_*NM*_ *> ϕ*_*M*_) and they reach a maximum for a volume fraction of the non-motile strain *ϕ*_*NM*_ ≃ 5 % and a volume fraction of the motile strain *ϕ*_*M*_ ≃ 0.2 % (Fig. 1c). The range of densities over which the fluctuations are observed corresponds to typical planktonic culture densities for the motile cells (0.01 *< ϕ*_*M*_ *<* 1 %) and typical biofilm densities for the non-motile ones (0.1 *< ϕ*_*NM*_ *<* 10 %), suggesting that this phenomenon is highly relevant to natural bacterial communities. Moreover, the heterogeneity level of the non-motile strain decreases at high volume fraction of motile cells (*ϕ*_*M*_ ≥ 1 %), where collective motion begins to occur ^26,29,30^. Interestingly, the spatial distribution of the motile strain was always homogeneous (Supplementary Fig. S3), and the density fluctuations only appear in the non-motile strain when the motile strain was present (Fig. 1b, d).

Therefore, the swimming activity of the motile cells drives the emergence of these out-of-equilibrium density fluctuations for the non-motile cells, without them being affected in any obvious way. Moreover, no change in the swimming speed was noticed. This is contrary to the behavior in simulated MIPS-like phase separation of mixtures ^32,33,52–57^, suggesting that a different mechanism is at play. To gain more insight into this mechanism, we systematically explored possible control parameters of the behavior.

### Tumbling, but not chemotaxis, affects the density fluctuations via the persistent circular swimming of surface localized motile cells

Since chemotaxis is known to be involved in the formation of some bacterial density patterns ^38–42,66,67^, we investigated whether this is the case in our system. We targeted the diffusible response regulator CheY, which plays the essential role of transducing through the cytoplasm the sensory signals from the chemoreceptors that monitor environmental conditions to the flagellar motor, where it triggers tumbles when phosphorylated ^10–12^. We first used a CheY knock-out (Δ*cheY*) motile strain, which is non-tumbling and non-chemotactic. To our surprise, the density fluctuations of the non-motile strain were found to be stronger and to appear on a wider range of cell densities in a mixture with the Δ*cheY* motile strain (Fig. 2a, b, c) than when the motile strain is wild-type (WT) for chemotaxis (Fig. 1c). Although the typical size of the density patterns is largely unaffected (Supplementary Figs. S1d,e), the local extrema of density are much more accentuated, with numerous depletion zones forming next to very dense patches (Fig. 2a, Supplementary Movie 3). However, a heterogeneity level similar to the WT strain case was found across the whole phase diagram for the motile *cheY* ** mutant (Supplementary Fig. S4a), for which a double point mutation (D13K, Y106W) in the *cheY* gene allows WT level of tumbling at proper expression level, but prevents chemotactic signaling ^68^. This indicates that chemotaxis *per se* is not important for the emergence of the density fluctuations, but that the ability to tumble is a control parameter of the phenomenon.

**Figure 2:**
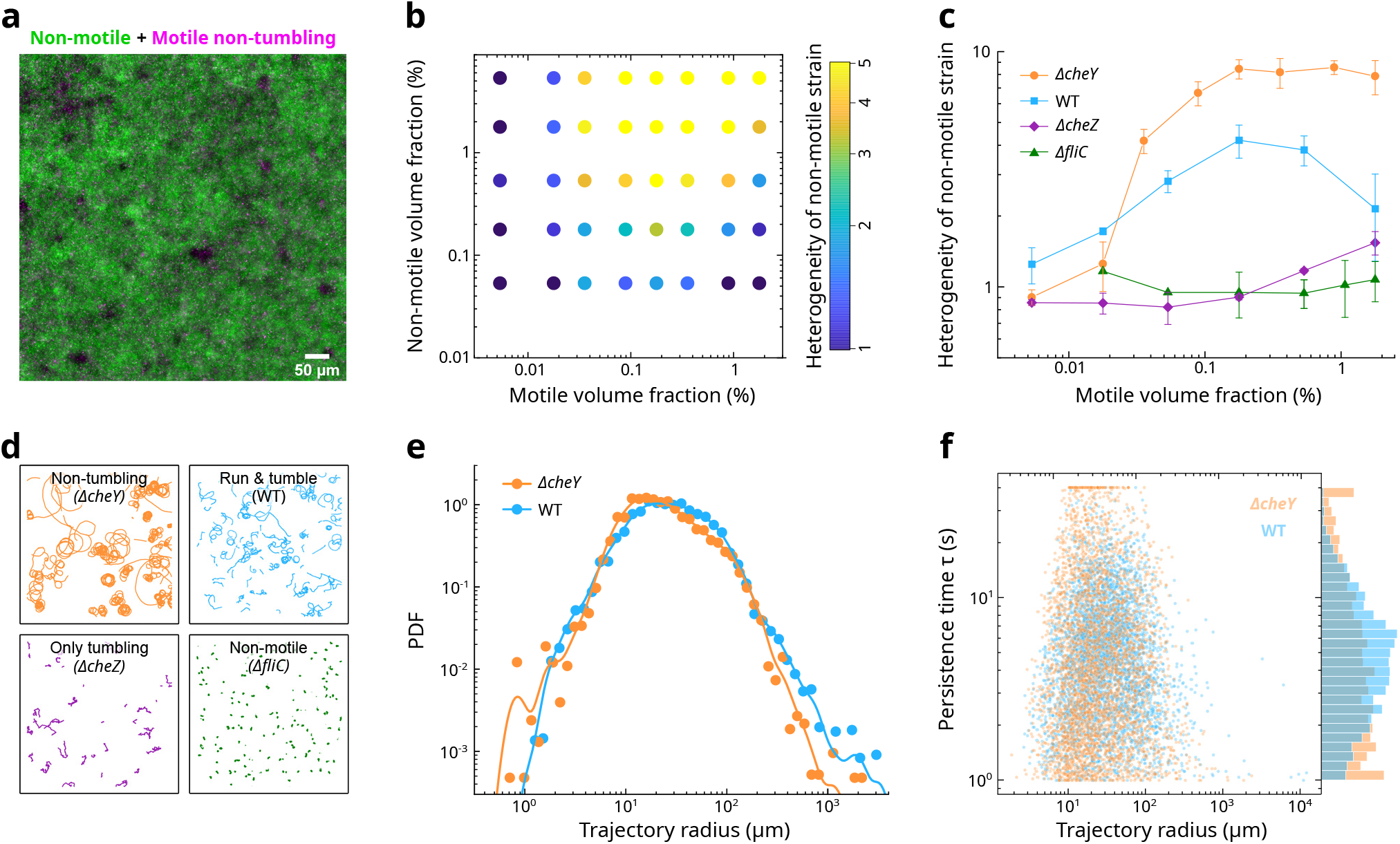
Tumbling affects the density fluctuations. a) Microscopy image of *E*.*coli* mixtures of a motile non-tumbling strain (Δ*cheY* mCherry) and a non-motile strain (mNeonGreen) where the non-motile strain presents large density fluctuations in the presence of the motile strain. b) Phase diagram of the density fluctuations in mixtures of a non-tumbling motile strain (Δ*cheY*) (volume fraction in abscissa) with a nonmotile strain (volume fraction in ordinate). Density fluctuations are quantified as described in Methods by the heterogeneity level, which is represented with the same color scale as Fig. 1c,d. c) Heterogeneity levels of the non-motile cells in a binary mixture with different motile strains as indicated, as a function of the volume fraction of the motile strain. The volume fraction of the non-motile strain is kept constant (*ϕ*_*NM*_ = 5.3%). (b,c) Each point represents the geometric mean of three replicates. (c) Error bars indicate the standard deviation. d) Swimming trajectories of the different strains used as the motile strain in the binary mixtures. e) Probability density distribution of the radii of curvature of individual circular swimming trajectories of WT and Δ*cheY* motile cells, which were tracked at the bottom of an experimental chamber. Each trajectory is weighted by its length. Lines represent the kernel density estimate of the data. f) Scatter plot of the persistence time *τ* of the same trajectories as (e), as a function of the radius of curvature of the trajectory, and probability density distribution of *τ* (right side) (e-f) The dataset comprises 4137 (Δ*cheY*) and 4475 (WT) individual swimming trajectories in 8 biological replicates.

From a physical standpoint, tumbling is known to help escape surface entrapment^22^, and thus to reduce the localization of motile cells to the surfaces ^19,20^. We measured the vertical distribution of motile cells in our system via confocal microscopy, and found that their preferential localization near the top and bottom surfaces was indeed enhanced in the Δ*cheY* strain compared to WT, but only to a moderate level (Supplementary Fig. S4b).

Moreover, close to the surfaces, the flagellated bacteria were observed to swim in clockwise circles when observed from above (Fig. 2d) as previously reported, because of the hydrodynamic torque induced by the no-slip wall ^23^. We measured the distribution of radii of curvature of the circular trajectories, which was very broad and log-normal both for the WT and non-tumbling strains (Fig. 2e). This broadness is not unexpected since the radius of curvature of the trajectory depends strongly on the variable distance between the cell and the surface ^23,24^. The distributions were almost identical for the WT and non-tumbling strains (Fig. 2e). However, in the Δ*cheY* strain, the circles are perturbed only by rotational diffusion. They thus tend to persist in place much longer than in the WT strain, where tumbles interrupt them (Fig. 2d). We quantified the time needed for a cell to “break the circle” by measuring a circle persistence time defined as the typical time *τ* each swimming cell takes to move a linear distance *dr*(*τ*) ≥ 3*R*, with *R* the mean radius of curvature of the trajectory. The distribution of *τ* is strongly shifted to higher *τ* for the Δ*cheY* strain compared to WT, with a large fraction of Δ*cheY* cells staying in the ≥ 3*R* vicinity for the whole duration (40 s) of our cell tracking movies (Fig. 2f). Persistent swimming in circles at the surface thus appears to be key for the emergence of the density fluctuations.

We next attempted to perturb the circular swimming of the motile cells. We first altered the distribution of radii of curvature by elongating both WT and Δ*cheY* motile cells via a short treatment with the antibiotic cephalexin ^29^. Elongated cell trajectories show an average increase of the radii of curvature, as expected from theory ^23^, but their distribution remains broad, likely due to the effect of the distance to the surface (Supplementary Fig. S5a-d). When mixed with elongated motile cells, the heterogeneity level of the non-motile cells is unchanged for Δ*cheY* and slightly increased for WT motile cells (Supplementary Fig. S5e). Because elongated cells also have a larger hydrodynamic dipole strength, and thus notably elicit larger hydrodynamic flows that could affect the non-motile cells, this result hints towards less curved trajectories being less potent in generating heterogeneities, all else being equal. We next drastically altered the swimming pattern by considering a tumbling-only motile strain, Δ*cheZ*. This strain’s flagella rotate, but only clockwise, causing constant tumbling and Brownian-like random walk (Fig. 2d), while still pouring energy into the fluid. The heterogeneity level of the non-motile strain of the mixture drops to almost control levels for all tumbling-only volume fractions (Fig. 2c), highlighting the importance of swimming patterns being spatiotemporally organized.

Taken together, these results indicate that the persistent and strongly curved circular swimming trajectories drive the formation of the density fluctuations.

### Sedimentation of non-motile cells is essential for the density fluctuations

We next considered the vertical distribution of non-motile cells, which is similar whether the motile strain of the mixture is WT or Δ*cheY* (Supplementary Fig. S4c). In both conditions, non-motile cells localize towards the bottom of the sample because of sedimentation, their volumetric mass density (*ρ*_*C*_ = 1.1 g.cm^*−*3^) being higher than the one of the aqueous medium (*ρ*_*w*_ = 1.0 g.cm^*−*3^). The (Boltzmann) steady state vertical distribution of sedimenting particles results from a balance between sedimentation and diffusion. The latter is enhanced as the volume fraction of motile cells increases (Supplementary Fig. S6a,b), as previously observed ^43,44,46^, and the vertical distribution of non-motile cells therefore becomes more homogeneous in presence of an increasing density of motile cells (Supplementary Fig. S6c,d).

Inspired by previous works on sedimentation of high-density colloidal suspensions ^69,70^, we decided to investigate how sedimentation affects the density fluctuations in our system. We matched the volumetric mass density of the suspending medium to the one of the bacteria by adding iodixanol, a non-ionic, non-cytotoxic iodine compound that is commonly used as a density matching agent and density gradient medium. It also has little effect on the suspension’s viscosity and is metabolically inert to *E. coli*. Coherently, adding iodixanol has little to no effect on the swimming speed of motile cells (Supplementary Fig. S7a,b) and on the diffusion of non-motile cells (Supplementary Fig. S7c). In the neutrally buoyant medium (Δ*ρg* = 0), the vertical distribution of non-motile cells becomes homogeneous, and thus top-down symmetric (Fig. 3a and Supplementary Fig. S7d), contrary to the asymmetric distribution in the negatively buoyant medium (Δ*ρg* = 1, in normalized units).

**Figure 3:**
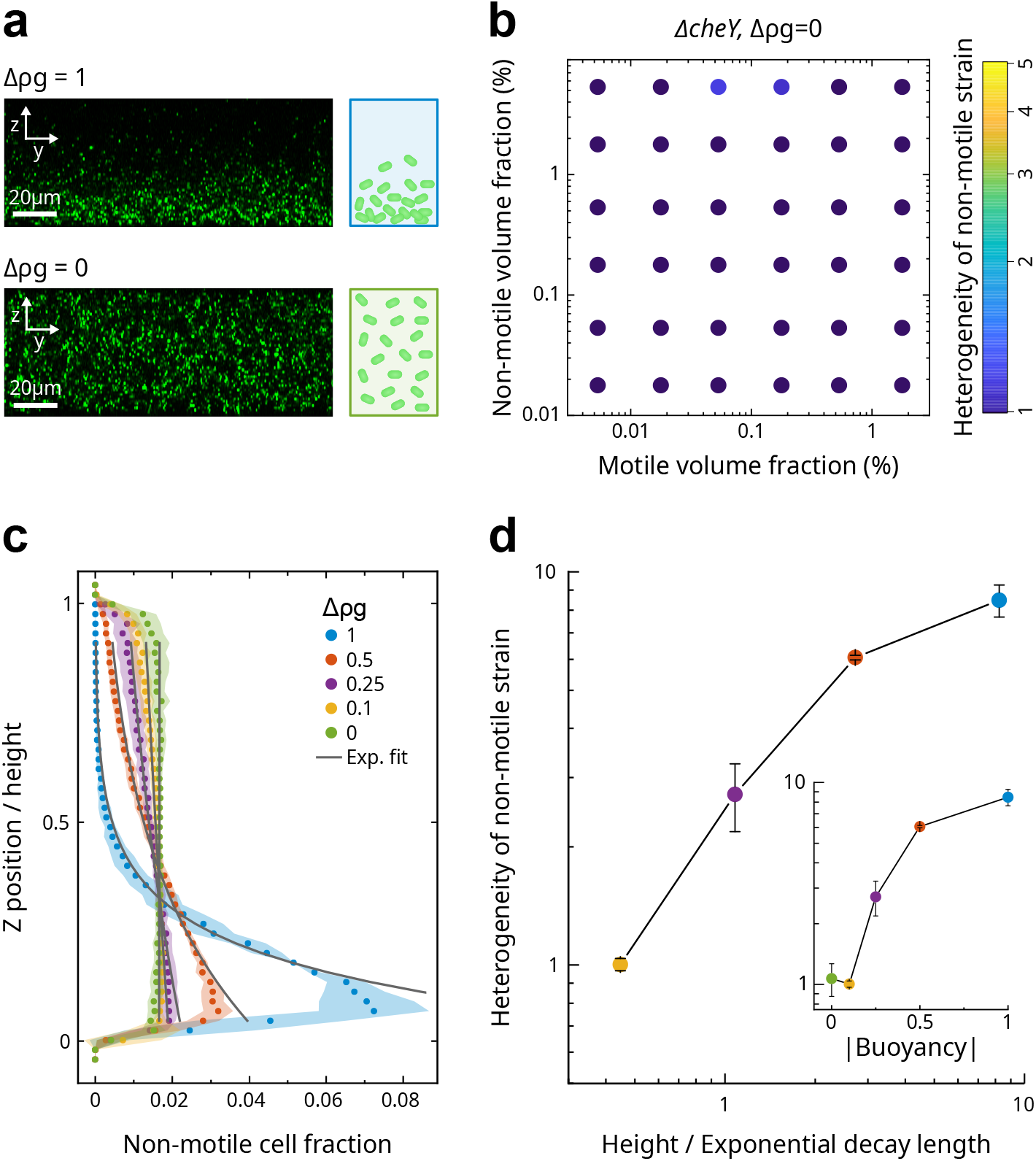
Sedimentation controls the density fluctuations. a) Confocal images of the vertical distribution of non-motile cells at the two extreme buoyancies Δ*ρg* = 1 and Δ*ρg* = 0. b) Phase diagram of the density fluctuations in a mixture of Δ*cheY* motile strain (volume fraction in abscissa) with a non-motile strain (volume fraction in ordinate) in density-matched suspending media (Δ*ρg* = 0). The density fluctuations are quantified by the heterogeneity level (Methods) which is represented by a color scale. c-d) Dose-dependence effect of buoyancy on the sedimentation (c) and the density fluctuations (d) of the most heterogeneous binary mixture (*ϕ*_*M*_ = 0.18%, *ϕ*_*NM*_ = 5.3%, motile strain Δ*cheY*). c) Non-motile cell fraction (abscissa), measured by cell counting in confocal microscopy, as a function of the vertical position (z) in a channel of 50 *µ*m height (ordinate), for suspensions at different buoyancies (the buoyancy level Δ*ρg* is normalized for motility buffer (Δ*ρg* = 1)). The grey lines are exponential fits of the cell fraction as a function of z. d) Density fluctuations of the non-motile strain as a function (main figure) of the inverse exponential decay length of the sedimentation profile and (inset) of the buoyancy normalized for motility buffer (Δ*ρg*). a-d) Each point represents the geometric mean of three replicates. Error areas (c) and bars (d) indicate the standard deviation.

In absence of sedimentation, the density fluctuations of the non-motile cells are completely annihilated, whether the motile strain in the mixture is tumbling (WT, Supplementary Fig. S7e) or not (Δ*cheY*, Fig. 3b). The level of heterogeneity is indeed equal to the one in the non-motile-only control (Fig. 1d). Focussing on one of the most heterogeneous mixtures when Δ*ρg* = 1 (motile Δ*cheY* with *ϕ*_*M*_ = 0.18% and *ϕ*_*NM*_ = 5.3%), we investigated the effect of varying the difference of volumetric mass density between the cells and the medium,

i.e. the sedimentation speed, and therefore the slope of the vertical cell density profile (Fig. 3c). The amplitude of the density fluctuations increased as a function of this density mismatch (Fig. 3d), showing that the cell sedimentation speed is a control parameter of the system.

Furthermore, no matter how high we increased the density of non-motile cells, we never observed any density fluctuations in the Δ*ρg* = 0 case (Supplementary Fig. S7f). We also confined the mixtures in a 20 *µ*m high channel, for which the half channel height matches the scale of the vertical gradient of non-motile cell density (Supplementary Fig. S8a). No density fluctuations were observed in this case (Supplementary Fig. S8b), further highlighting the importance of the three-dimensionality of the system. These results strongly indicate that the effect of sedimentation is not merely due to an increased density of cells near the bottom surface, but that the vertical gradient of cell density resulting from sinking under gravity is essential for the formation of the density fluctuations.

We then studied the dynamics within the density fluctuations at steady state (Supplementary Movie 1 & 3). The non-motile cells show collective movements with many converging and diverging convective flows, somewhat resembling boiling water. The movements were more pronounced in presence of non-tumbling motile cells, with cell depletion zones (“holes”) intermittently opening and closing. The horizontal cell motion was quantified using particle image velocimetry, with the non-motile cells directly serving as tracers (Fig. 4a, b). Even at volume fractions of motile cells far below the emergence of bacterial turbulence, the velocity fields revealed large spatial correlations in the motion of non-motile cells (Fig. 4a). To quantify these correlations, the normalized spatial power spectral density of the velocity field, 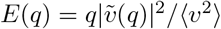, was computed as a function of the spatial wave number *q* (Methods) ^28,29^. It represents the fraction of the kinetic energy (*v*^2^) in flow structures of typical size *π/q*. E(q) shows a pronounced peak, the hallmark of spatial correlation, at a wavelength *π/q* = 50 *µ*m, corresponding to the height of the channel (Fig. 4b). This value of the correlation length indicates that hydrodynamic interactions drive these collective movements of the non-motile cells, similarly to what was observed for swirling collective motion of homogenous populations of swimmers ^29^ and suggests that the motile cells generate structured flow fields that are able to stir the non-motile ones. The temporal correlations of the velocity field decayed rapidly in the tumbling motile case, whereas long term correlations are observed in the non-tumbling one (Supplementary Fig. S9a), coherently with the longer persistence of circular swimming in the latter case (Fig. 2f).

**Figure 4:**
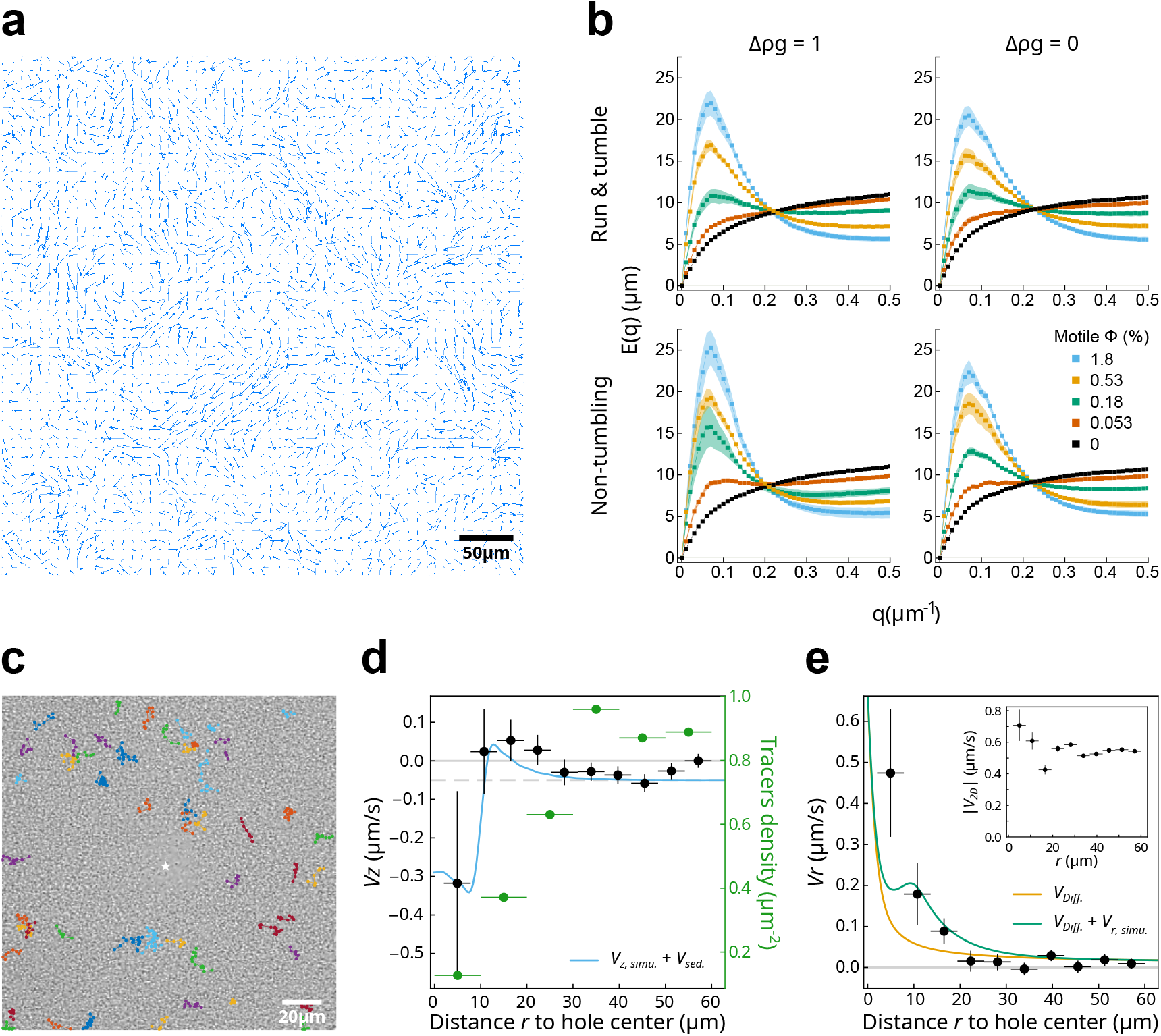
Dynamics of the non-motile cells in the density fluctuations. a) Instantaneous velocity field of the non-motile strain at *ϕ*_*NM*_ = 5.3% in mixture with the motile strain Δ*cheY* at *ϕ*_*M*_ = 0.53% and at buoyancy Δ*ρg* = 1. b) Flow structure factor E(q) of the non-motile strain flow field for increasing cell densities and for mixtures with the indicated motile strains and buoyancies. c) Field example of the three-dimensional tracking of non-motile cells close to a hole performed in the mixture *ϕ*_*NM*_ = 5.3%, Δ*cheY ϕ*_*M*_ = 0.53%, at buoyancy Δ*ρg* = 1. The white star represents the mean position of the hole center over the whole recording time, and the colored points represent the XY projection of the 3D trajectories of the tracers (tagged nonmotile cells). d) Average vertical velocity of the tracers (*V*_*z*_) (black circles) and their density (green circles) as a function of their distance *r* to a hole. e) Average radial velocity of the tracers (*V*_*r*_) (black circles) and horizontal speed (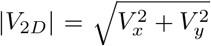, inset) as function of the distance *r* to a hole. d-e) The velocities were quantified from 639 individual 3D trajectories situated less than 60 *µ*m away from a hole center. Each data point represents the binned mean and the vertical error bars represent the standard error. The continuous lines are simulated flows generated by circular swimmers (*R* = 10 *µ*m, *a*^*S*^*v*_0_ = 70 *µ*m^2^/s). (d) The vertical velocity sums the flow of the circular trajectories (*V*_*Z,simu*._) and the sedimentation (*V*_*sed*_ = 0.05 *µ*m) (blue line), (e) and the horizontal velocity result from the diffusive flux (*V*_*Diff*._) alone (yellow line) and with the effect of the circular trajectories (*V*_*r,simu*._) (green line).

The level of heterogeneity is however not well predicted by the strength of the collective flows alone, as measured by the height of the peak of *E*(*q*) (Supplementary Fig. S9b). Moreover, a peak of *E*(*q*) was also observed in the density-matched suspending medium, which appears at the same length scale and with barely lower amplitude than the one in the corresponding negative buoyancy case (Fig. 4b), indicating quasi-identical spatial flow structures whether Δ*ρg* = 0 or Δ*ρg* = 1. However, the total amount of kinetic energy (*v*^2^) is somewhat lower in the neutral buoyancy case (Fig. S9c). Hence, regardless of the buoyancy, the motile cells are found to generate spatially correlated movements of the non-motile cells via hydrodynamic interactions, which however only give rise to strong heterogeneities of density in the negative buoyancy case, when the vertical symmetry of the non-motile cell density is broken.

We thus next considered the vertical motion of the non-motile cells. We tracked the three-dimensional motion of a few marked non-motile tracers with confocal microscopy in mixtures with unmarked non-motile and non-tumbling motile cells. We focused on areas where cell depletion zones (“holes”) are visible with transmitted light, in the Δ*ρg* = 1 case (Fig. 4c), and quantified the velocity of the tracers as a function of the distance to the center of a hole (*r*) (Fig. 4d,e and Supplementary Fig. S9d,e). Close to the hole (*r <* 10 *µ*m), the mean vertical velocity of the tracers (*V*_*z*_) is large and negative, i.e they move down fast on average, while they move on average rapidly away from the hole in the horizontal direction (*V*_*r*_ *>* 0). Further away from the hole (*r* ∼ 20 *µ*m), the mean vertical velocity reaches a maximum, and the typical horizontal motion (|*V*_2*D*_|) marks a lull. Far from the hole (*r >* 30 *µ*m), the radial velocity relative to the hole (*V*_*r*_) is zero on average, while the mean vertical velocity is consistent with the sedimentation speed of the non-motile cells (≃ 0.05 *µ*m/s). These findings suggest that the non-motile cells are exposed to a recirculatory flow, being dragged down and expelled away from the hole when close to it.

### Recirculative flows generated by circular swimming generate density fluctuations in simulations

We turned to simulations to understand how the fluid flows generated by cells swimming in circle close to the surface allow for density fluctuations to emerge. We used a *N* log *N* algorithm ^71,72^ to compute fluid flows under confinement by the two no-slip parallel walls at the top and bottom of the simulation domain, with periodic boundary conditions in the other directions (Methods and Supplementary Notes). The swimming cells are approximated as force dipoles, with a dipole length *l*_*p*_ = 4 *µ*m, and a reduced force *f*_0_*/*6*πη* = *a*^*S*^*v*_0_, with *a*^*S*^ the swimmer hydrodynamic radius and *v*_0_ the swimming speed. We first computed the normalized flow field generated by a single dipole that describes a perfectly circular trajectory, averaged over one circle revolution, *v*_1*r*_(*r, z*)*/a*^*S*^*v*_0_ (Fig. 5a). The flow field is axisymmetric around the center of the circle, and features a vertical recirculation, where the fluid is sucked down in the middle of the circle to be expelled out and up on its outside (Fig. 5a and Supplementary Fig. S10a-f). The curvature of the dipole trajectory sets the strength of the recirculation, with the magnitude of the typical fluid velocity scaling as ⟨|*v*_1*r*_|⟩_*r,z*_ ∝ 1*/R*^3^ with the recirculation radius *R* (Methods and Supplementary Fig. S10g-i), while the maximal velocity scaled as max ⟨|*v*_1*r*_|⟩ ∝ 1*/R*^2^ (Fig. 5b). Consistently, the recirculation flux (equation 12 and Methods), was found to be proportional to 1*/R* (Fig. 5c). The curvature of the dipole trajectory thus creates a net outward force perpendicular to the circle that drives the recirculation.

**Figure 5:**
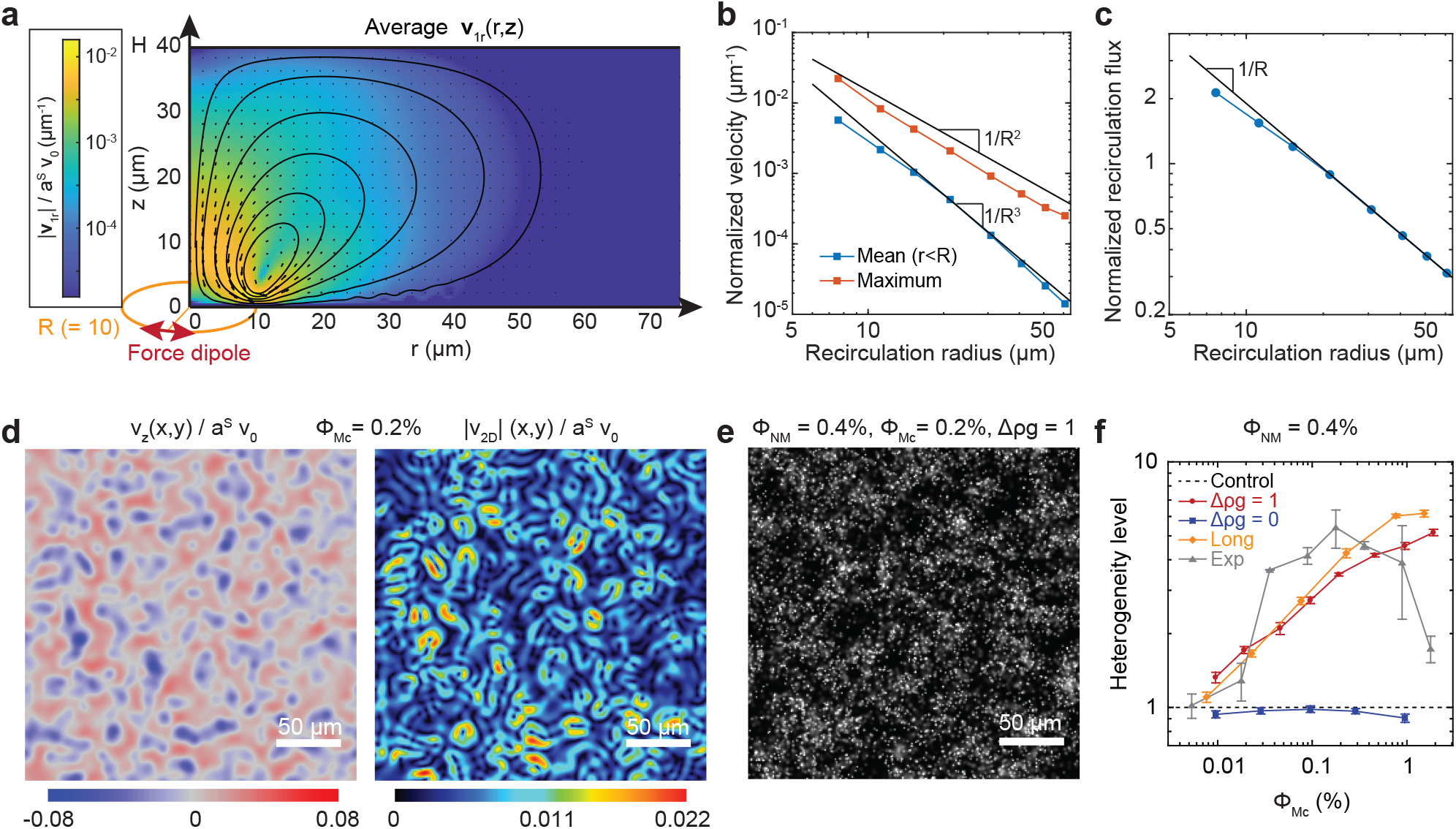
Recirculation flows lead to heterogeneities of non-motile cells in simulations of circular swimmers at surfaces. a, Map of the axisymmetric mean flow field generated by a circularly swimming force dipole. The flow field is computed between two no-slip walls spaced by *H* = 40 *µ*m, and the dipole circulates *z*_*M*_ = 1 *µ*m (about one cell body) above the bottom wall. Shown are the flow speed normalized to dipole force *a*^*S*^*v*_0_ (color), the velocity vectors (arrows) and streamlines in the (r,z) plane (solid lines), featuring a recirculation around a fixed point close to the circle radius. b, Mean and maximum of the flow speed inside the circle as a function of the position of the fixed point (recirculation radius). Solid lines indicate power law dependences. c, Normalized recirculation flux, max_*{r}*_ Φ(*r*)*/a*^*S*^*v*_0_ (equation 12, in *µ*m) as a function of the recirculation radius. The solid line indicates 1*/R* dependence. d, Example of mean flow field generated by a population of circular surface swimming dipoles (density *ϕ*_*Mc*_ = 0.2 %) for the distribution of circle radii of Fig. 2f. The vertical (*v*_*z*_) and horizontal 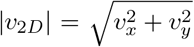 components are shown in the (x, y) plane, *z* = 8 *µ*m above the bottom wall. Color scales indicate *v/a*^*S*^*v*_0_ in *µ*m^*−*1^. e, Reconstructed microscopy image of negatively buoyant diffusive tracers (*v*_*sed*_ = 0.05 *µ*m/s, *D* = 0.3 *µ*m^2^/s) that followed the mean circular swimmers flow field for 100 seconds, for the same conditions as (d). f, Heterogeneity level of the non-motile tracers, computed in simulations as a function of the volume fraction of motile circular swimmers *ϕ*_*Mc*_ for non-motile cells (NM) with volume fraction *ϕ*_*NM*_ = 0.4%. Control: Homogeneous distribution, Δ*ρg* = 1: sedimenting NM, Δ*ρg* = 1: non-sedimenting NM, Long: elongated motile cells. Exp: experimental data for non-tumbling motile cells, Δ*ρg* = 1 and *ϕ*_*NM*_ = 0.53%. Error bars for simulated data indicate standard error of the mean over 6 repeats. Experimental data encompass 3 biological replicates.

The shape of the simulated flow curves matched well with our experimental measurements close to depletion zones (Fig. 4d,e and Supplementary Fig. S10b,d,f). We vertically averaged *v*_1*r*_(*r, z*)*/a*^*S*^*v*_0_ over the experimental distribution of tracer’s z-positions in the 3D-tracking experiments, for a circle radius *R* = 10 *µ*m, which corresponds well to the apparent size of the typical experimentally observed depletion zone. This average simulated velocity field was in good agreement with the experimental velocity field around a depletion zone, when sedimentation and diffusion away from the hole center are also taken into account (Fig. 4d,e). The only fit parameter was the reduced dipole force, *a*^*S*^*v*_0_ = 70*µ*m^2^/s, which is 5 times larger than the expected value *a*^*S*^*v*_0_ ∼ 14*µ*m/s (*a*^*S*^ = 0.7 *µ*m and *v*_0_ = 20 *µ*m/s as measured for our motile cells). This discrepancy might come from the steep dependence of *v*_1*r*_ on the radius *R* and/or from several circular swimmers contributing to various degrees to the hole formation.

The vertically recirculating flow field might thus perturb the bottom–heavy distribution of passive cells to form depletion zones surrounded by higher densities as observed experimentally. To understand how this effect translates for thousands of interlaced circular swimmers, we simulated the mean flow field generated by a population of dipoles that swim in circle, with an experimentally-grounded distribution of circle radii (Supplementary Fig. S11a and Methods). The trajectory centers are evenly distributed at both surfaces and independent from each other. Even for large swimmer densities (e.g. *ϕ*_*Mc*_ = 0.2%, *n*_*Mc*_ = 5 10^3^ circles, for which the experimental heterogeneity level is maximum), the normalized total flow field *v*_*tot*_*/*(*a*^*S*^*v*_0_) shows a few prominent recirculation patterns over a more noisy background (Fig. 5d). Indeed, the typical contribution of a circle of radius *R*_*i*_ being of order 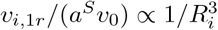, even for very large numbers of swimmers, the relatively few smaller circles in the distribution contribute massively to the total mean flow field and make it spatially heterogeneous (Fig. 5b and Supplementary Fig. S11b). The flow speed, using *a*^*S*^ = 0.7 *µ*m and *v*_0_ = 20 *µ*m/s, are of order *v*_*z*_ ∼ 0.5 − 1 *µ*m/s in the fast areas, which is now compatible with our experimental measurements (Fig. 4d,e), and suggests that heterogeneities arise from the flow field.

We thus computed the trajectories of passive tracers, which experienced gravity (*v*_*sed*_ = 0.05 *µ*m/s) and Brownian motion (*D* = 0.3 *µ*m^2^/s) at the level of non-motile cells in Δ*ρg* = 1 conditions, in the mean flow field generated by fixed configurations of circular swimmers with *a*^*S*^*v*_0_ = 14 *µ*m^2^/s. For motile and non-motile densities in the experimental range, we observed the formation of strong patterns of density fluctuation that are similar to the experimental ones on reconstructed images of non-motile cells (Fig. 5e). They show giant number fluctuations (Supplementary Fig. S11c), which we used to define a level of heterogeneity (Methods), comparably to the experiments (Supplementary Fig. S1). The level of heterogeneity increases as a function of the time *τ* during which the tracers are exposed to a fixed configuration, corresponding in experiments to the time during which circles would persist (Supplementary Fig. S11d). For the highest fractions of circular swimmers, we also observed a partial vertical homogenization of the density distribution of non-motile cells (Supplementary Fig. S11e), similarly to experiments (Supplementary Fig. S6c,d). For a persistence time of *τ* = 60 s, which is consistent with the experimental non-tumbling swimmer case (Fig. 2f and Supplementary Fig. S9a), the dependence of the simulated heterogeneity level on circular swimmer density is in fairly good quantitative agreement with experiments (Fig. 5f and Supplementary Fig. S11f), modulo a purely geometric scaling factor (Methods). If gravity is set to zero, no heterogeneities are observed in simulations, in agreement with experiments again, because non-motile cells are homogeneously distributed and the flows are incompressible (Fig. 5f). Normal and elongated circular swimmers also elicit equal levels of non-motile heterogeneities (Fig. 5f) in agreement with our experiments with the non-tumbling motile strain (Supplementary Fig. S5). Our simulations therefore recapitulate our experimental results and provide a mechanism for the emergence of the density fluctuations in binary mixtures: Circular swimming trajectories close to surfaces generate a vertical recirculation that disturbs the asymmetric distribution of sedimenting non-motile cells.

## Discussion

In this paper, we report the emergence of density fluctuations in binary mixtures of motile and non-motile bacteria, over a broad range of physiologically relevant volume fractions of both species, and we identified the purely physical mechanism of this emergence, based on the combination of heterogeneous active flow fields with a symmetry breaking by gravity. Our experiments and modeling indeed demonstrated that persistent swimming in circles of the motile cells at the surface of the observation chamber, where they accumulate, creates recirculation fluid flows with regions of downwards suction surrounded by upward lift of the fluid, which collectively convect the non-motile species in three dimensions. At the population level, these flows are heterogeneous because the distribution of radii of curvature *R* of the swimming trajectories is broad and the typical fluid velocity produced by a circular swimmer scales as 1*/R*^3^, resulting in infrequent small *R* trajectories strongly contributing to the total flow. The second ingredient for the active density fluctuations to emergence is sedimentation under gravity, which breaks the vertical symmetry of the system, and allows cells to accumulate in the regions of upward lift, while they are depleted from the suction regions. In contrast, the fluid incompressibility prevents density heterogeneities to emerge in absence of gravity. Finally, this mechanism is strongly supported by numerical simulations of sedimenting non-motile particles in the flow generated by populations of surface circular swimmers, which could recapitulate all our experimental observations.

Our experiments clearly establish the sedimentation inverse length (1*/z*_*sed*_ = *v*_*sed*_*/D*_*NM*_) as one of the control parameters of the system (Fig. 3). The decrease of the heterogeneity level of the non-motile cells at high motile cell density can then be understood as a consequence of the co-occurrent increase in the effective diffusion coefficient of non-motile cells, which reduces 1*/z*_*s*_ and homogenizes them in the vertical direction (Supplementary Fig. S6). Our simulations and experiments agreed very well despite our rather strong approximations. Steric interactions become important at the non-motile densities we considered. Including them might thus improve this agreement further. Collective sedimentation effects, which can produce heterogeneities by themselves ^69,70^, are also likely to reinforce the heterogeneities produced by the motile cells and might explain the somewhat lower heterogeneity level in simulations compared to experiments (Fig. 5f). Our model also assumes that the swimmer force dipoles are parallel to the surface, although evidences suggest that the flagellum sticks up with an angle at the surface ^18,21^. The dipole flow might then be slightly asymmetric, with a stronger contribution of the flagellum. The implied additional azimuthal component to the recirculation flow field was however not visible in our experiments, suggesting that such an effect is small at best here. Our simulations, as well as experiments, indicate that the persistence time of the circular swimming controls the strength of the heterogeneities (Fig. 2f). This persistence time must decrease as a function of tumbling rate, tumbling angle and rotational diffusion ^20,22,24,25^. The latter two are reduced in elongated cells ^73^, which probably contributes to the increased heterogeneity level upon elongation of WT cells (Supplementary Fig. S5). Interaction with the surface are furthermore known to affect tumbling ^74^ and possibly diffusion ^25^. Therefore, this interaction might also reinforce the recirculative flows by allowing some long persistence of the circular motion in the WT case.

The emergent active density fluctuations we uncovered constitute a new type of collective structuration for bacteria. These density fluctuations indeed differ from MIPS-based phenomena ^32–36,57,58^, given that the spatial localization of the motile cells is not affected (Supplementary Fig. S3) and their influence on the nonmotile cells can be explained only considering hydrodynamic interactions (Fig. 4). Our results thus highlight the importance of hydrodynamic interactions even below the onset of swirling collective motion known as bacterial turbulence ^26,29–31^. In the most heterogeneous mixtures, the fluid flows are not strong enough to visibly affect other swimmers, but a flow structure has already emerged, which is revealed by the non-motile cells. This flow structure might have other signatures, which were up-to-now unrecognized, for example previously observed transient enhancement of chemotactic drifts ^29^. Although our system requires three-dimensionality (Fig. S8), the behavior we observe shares similarities with FDPO, which is conceptualized on fluctuating surfaces ^61–63^, including giant number fluctuations of cell density and non exponentially decaying correlation functions (Supplementary Fig. S1). The temporally varying space-structured flow field created by the swimmers, together with the gravity constraint, might then be interpretable in terms of a fluctuating surface ^61^. Experiments with rapidly sedimenting or two-dimensionally constrained beads have shown that swimming bacteria can generate attractive interactions between them ^47^, which seem to lead to collective aggregation ^48,49^. While these interactions were observed in two dimensions and mainly interpreted as active depletion interaction effects, the shape of the fluid flow generated by surface swimmers may also contribute to the observed behavior, in a low diffusion and infinite sedimentation limit of our experiments.

Active particles attached to surfaces can affect local dynamics, a phenomenon coined active carpet ^75^, by enhancing diffusion and creating fluid transport ^76,77^, mostly observed in eukaryotic cells like in the synchronously beating cilia ^78,79^ but also in such prokaryotic systems as engineered surfaces covered with attached flagellated bacteria ^80^. Our observations fit in this general framework, further highlighting the importance of microorganism-surface interactions. However, in our case, the structure emerges from a population of freely swimming cells interacting with the surface.

The results presented in this study give insight into the potential impact of motility on microbial community dynamics in natural settings. Given the magnitude of the flow fields generated by the swimming *E. coli*, non-motile but also slowly moving species (at speeds ≲ 1*µ*m/s) should be affected by the density fluctuations. We expect the phenomenon we uncovered to play a role in the enhancement of self-aggregation ^66,81^, and also to affect the formation of biofilms in complex multispecies communities. Further investigation is required to determine whether the structure generated by motile cells provides any hindrance or benefits to non-motile cells. Additionally, while chemotaxis is not necessary for the emergence of this phenomenon, it is likely that interactions via chemical cues between the community members modulate the behavior in more natural settings ^38–42,64–66^. Overall, these results shed light on the complex interplay between individual cells and their environment, and contribute to our understanding of how microorganisms interact and organize in nature.

## Methods

### Strains and plasmids

All strains used in this study are derived from *Escherichia coli* wild-type strain W3110 (RpoS+), for which the gene *flu*, which encodes for the major adhesin in our growth conditions, Antigen 43, was deleted ^38^. Gene deletions are carried out using the plasmid pSIJ8, which carries the lambda Red recombineering genes inducible with arabinose and the flippase recombinase gene inducible with rhamnose ^82^. The target gene is substituted with a PCR product that carries the Kanamycin (Kan) resistance cassette with lambda Red recombinase by adding 0.15% of arabinose. Recombinant cells are selected with the Kan resistance and then the marker is removed with the flippase recombinase by adding 50mM L-rhamnose to the culture. The deletion of the gene is checked by colony PCR and then the plasmid pSIJ8 is removed by growing the cells at 42°C. The non-motile strains lack the flagellin *fliC*. The chemotaxis response regulator *cheY* was deleted in the non-tumbling strain, and the phosphatase *cheZ* was deleted in the tumbling-only strain.

All strains are fluorescently labeled with fluorescent proteins mCherry or mNeonGreen expressed from a plasmid with pTRc99a backbone with a strong ribosome binding site (RBS2) ^83^. Motile strains were usually tagged with mCherry and non-motile strains with mNeonGreen. Reversing the tags gave identical results (Supplementary Fig. S1c). Growth medium is supplied with 100 *µ*M Isopropyl *β*-d-1-thiogalactopyranoside (IPTG) for fluorescent protein expression induction and 0.1 mg/mL Ampicillin as antibiotic resistance marker. The non-chemotactic but tumbling strain expresses *cheY* ^*D*13*K,Y* 106*W*^ (*cheY* **) from a plasmid with pTRc99a backbone transformed in the Δ*cheY* strain. No IPTG was added, the expression of CheY** resulted only from the leaky expression of the plasmid. This strain was tagged with mCherry expressed from pOSK237, a plasmid with pBAD backbone, adding 0.004% L-arabinose for protein induction and 50 *µ*g/mL Kanamycin as antibiotic resistance marker in the growth medium. All the strains and plasmids used in this study are listed in Table S1.

### Microfabrication

Molds were fabricated using standard photolithography and microfabrication techniques. The SU8 photoresist (Microchem^*TM*^) was spin-coated on a silicone wafer, prebaked, shaped by patterned UV exposure, postbaked and washed according to manufacturer instruction. The cast was then passivated with silane and checked using a profilometer. Poly-di-methylsiloxane (PDMS) was poured on the cast in a 1:10 crosslinker to base ratio, degassed, baked overnight at 65 °C, peeled off, cut to shape and covalently bound on isopropanol-rinsed microscopy high precision glass slides after oxygen plasma treatment. PDMS to glass covalent bonds were allowed to form for 20 minutes at room temperature and the devices were then filled with the samples.

### Phase diagram assays

#### Cell growth and sample preparation

Cells are grown overnight in liquid tryptone broth (TB) medium (1% Bacto tryptone + 0.5% NaCl) with appropriate antibiotics at 37 °C. The overnight culture is diluted 1/100 in 10 mL fresh TB supplemented with antibiotics and inducers and grown at 34 °C under shaking at 270 rpm for about 4h, until the optical density at 600 nm of the suspension reached OD_600_ ≃ 0.5.

For elongated cells, 100*µ*g/mL of cephalexin, which blocks cell division, are added 60 minutes (i.e. about 1.5 cell cycles) before harvesting the cells, yielding cells about 2-4 times as long as without treatment.

The whole culture is then centrifuged at 3.5krpm for 5 minutes to prevent any damage to the flagella of motile cells. The pellet is resuspended in motility buffer (MB) (10mM KPO_4_ + 0.1mM EDTA + 67mM NaCl + 0.01% Tween 80, pH=7.0) + 1% Glucose. This buffer allows cells to swim but not to grow. Tween 80 is a surfactant that prevents cell-surface adhesion ^84,85^. Cells are brought to a high density (OD_600_=60) by a subsequent centrifugation (3.5krpm, 5 minutes) and then 6 serial dilutions are done for each of the strains in MB. Each dilution of each strain is mixed with a dilution of the other strain in a proportion 1:1, to a total of 36 binary mixtures. Each mixture is then loaded in a chamber of a previously fabricated microfluidic device. The chamber consists of an inlet connected to an outlet by a straight channel of 50 *µ*m height, 1 mm width and 1 cm length. The channel is then sealed with grease to prevent fluid flows. For steady state measurements, images are acquired after letting the mixtures settle in the microfluidic device for at least 40 minutes. For time dependent measurements, only two channels are filled at the same time, and brought immediately under the microscope. Each condition was probed in at least three independent biological replicates.

#### Physically modified media

When varying the buoyancy of the medium, cells are grown and processed as described above, except that the pellet of cells is resuspended in density matching motility buffer after the last centrifugation and the serial dilutions are made in this same density matching medium. The latter is prepared using iodixanol to increase the density of the MB up to the same density of a single *E*.*coli* cell (1.11g/ml). The density matching motility buffer contains 26.6% iodixanol, the concentration of the other components being unchanged (10mM KPO_4_ + 0.1 mM EDTA + 67mM NaCl + 0.01% Tween 80 + 26.6% iodixanol). For the experiments where different buoyancies were tested, the volume fractions of iodixanol were: 0% (Δ*ρg* = 1), 13.2% (Δ*ρg* = 0.5), 19.9% (Δ*ρg* = 0.25), 23.9% (Δ*ρg* = 0.1), 26.6% (Δ*ρg* = 0).

#### Epifluorescence Microscopy

The density fluctuations are visualized using the Nikon Ti-E inverted fluorescence widefield microscope with a 20x (NA 0.75) objective and the mCherry (excitation filter 572/25, emission 645/90) and GFP (excitation filter 470/40, emission 525/50) filters. Images are acquired in the focal plane 9 *µ*m above the bottom of the microfluidic channel. Focal plane is maintained using the Nikon perfect focus system.

#### Density fluctuations quantification

A stack of images of 9 different positions spanning across the microfluidic channel is acquired 7 times for each fluorescence channel with the Andor Zyla sCMOS Camera and NIS software (1 px = 0.32 *µ*m, field of view = 2048 × 2048 px^2^, 50ms exposure). Further image processing is realized with Fiji (https://fiji.sc/). Each image in the stack is corrected for inhomogeneous illumination. For this, an illumination background image is computed at each cell density as the median fluorescence of a large set of images taken at this density. The intensity of this background image is then normalized by its mean. Each image of the stack is divided by the illumination background image corresponding to its cell density. To further correct for small remaining systematic biases in the intensity profile of each image, a quadratic polynomial filter is applied.

The spatial autocorrelation function of the corrected intensity is computed as:

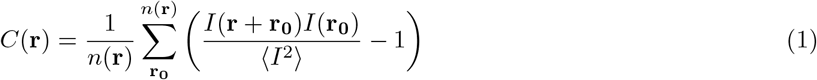

where *n*(**r**) is the number of valid initial pixel positions **r**_**0**_ for a given spatial shift **r**. In homogeneous control, this function decays over a length scale of about 1 *µ*m corresponding to the bacterial size, whereas a second decay appears at several tens of microns when density fluctuations are present (Supplementary Fig. S1e).

The density fluctuations are quantified by calculating the heterogeneity level of the images. For this, each image is divided into non-overlapping squares of size *N*, allowed to vary from *N* = (2^0^)^2^ px^2^ to *N* = (2^9^)^2^ px^2^ (Supplementary Fig. S1a). The sum of the fluorescence intensity 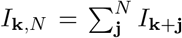 is computed within each square (indexed by **k**). The spatial variance 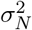 of this total intensity is then computed across the squares of size *N*,

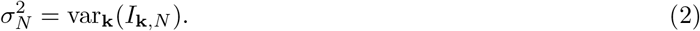

The latter is then normalized by the pixel size of the slice (*N*) and by the variance over squares of size *N* = 1, 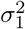 (Supplementary Fig. S1b), since one expects from the central limit theorem that 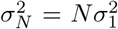 for an image made of pixels that assume random non-correlated values. The ratio 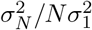 increasing as a power of *N* is then a marker of large scale correlations in the spatial distribution of fluorescence intensity, often called giant number fluctuations ^86^.

Since our images have a texture that comes from the cells being several pixels large, even in homogeneous samples, 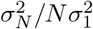 transiently increases with *N* until the latter reaches the area of a single cell (*N* ∼ 100 px^2^), above which 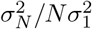 becomes constant as a function of *N*. In contrast, in samples with density fluctuations, 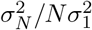 keeps increasing for *N >* 100 px^2^, marking the large scale fluctuations of the cell density (Supplementary Fig. S1b). To reduce the corresponding heterogeneity level to a single number, we use 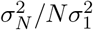 for a square of area *N* = (2^7^)^2^ px^2^ = 16384 px^2^. The width of this square is indeed about 42 *µ*m, which corresponds to the large scale decay of the spatial autocorrelation function of the pixel intensity (Supplementary Fig. S1e). Since the value of 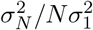 *for N* = (2^7^)^2^ in the control passive systems varies little across non-motile volume fractions and is close to 400 on average, we further defined the heterogeneity level as 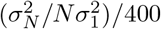 for *N* = (2^7^)^2^, such that passive systems have a heterogeneity level of 1 on average, and heterogeneity is defined relative to this homogeneous case.

#### Image velocimetry analysis

A movie of 7500 binned frames is acquired at 5 frames per second with the Andor Zyla sCMOS Camera and NIS software (1 px = 1.3 *µ*m, field of view = 512 × 512 px^2^, 50ms exposure). The velocity field, the flow structure factor and the kinetic energy of the non-motile cells are measured using a local image velocimetry algorithm based on Fourier image analysis, following previously described protocol ^29^ (Github code: 10.5281/zenodo.3516258). In short, local velocities are computed on a rectangular grid with 4 px spacing, using Gaussian filtered (half width 3 px) squared subframes (32 × 32 px^2^), with a 10 frames long sliding window. The spatial Fourier transform of the velocity field is computed as

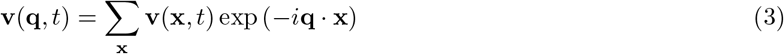

and the power spectrum as

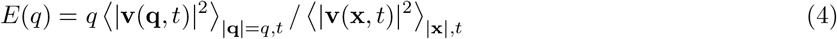

### Confocal microscopy

The binary mixtures are prepared following the above-mentioned Sample preparation and are visualized using a Zeiss LSM-880 confocal microscope equipped with a C-Apochromat 40x (NA 1.2) water immersion objective. Motile cells (tagged with mCherry) and non-motile cells (tagged with mNeonGreen) are visualized using the 594nm Helium-Neon laser and 488nm Argon laser, and detected with a photomultiplier at 490-535nm and 598-660nm, respectively.

### Cell fraction vertical distribution

Z-stack images of the whole microfluidic channel are acquired using the ZEN Black software (Zeiss) (1px = 0.2 *µ*m in X and Y, 1px = 1 *µ*m in Z; field of view = 303.64 × 303.64 × 70 *µ*m^3^, 0.35 *µ*s/px exposure). The number of cells in each Z plane is quantified by the connected components labeling algorithm for ImageJ ^87^. Because the height and the tilt of the microfluidic channels slightly vary from sample to sample (*<* 10%), the Z position is binned and the mean of the cell fraction over the bins is calculated.

### 3D tracking

To track the horizontal and vertical motion of the non-motile cells, the binary mixtures are prepared following the above-mentioned Sample preparation. To be able to track single non-motile cells, only the 0.03% of them are fluorescently labeled. Low density regions (“holes”) of the mixtures are identified with transmitted light and Z-stacks are acquired at the highest possible speed around these regions using the ZEN Black software (Zeiss) (1px = 0.54 *µ*m in X and Y, 1px = 2 *µ*m in Z; field of view = 163.5 × 163.5 × 20 *µ*m^3^, 1.3 *µ*s/px exposure, ∼0.3fps).

The three-dimensional trajectories are reconstructed using the Mosaic plugin for ImageJ^88^. From each trajectory is then extracted their relative position to the hole center (defined by transmitted light) and their velocities in the vertical and horizontal planes.

### Motility assay and circular trajectories measurements

From an overnight culture, cells are grown in TB for 4h at 34°C. The culture is centrifuged (3.5krpm for 5 minutes), and the pellet is resuspended in MB + 1% Glucose. Cells are centrifuged again (3.5krpm for 5 minutes) and resuspended in the adequate buffer (MB, MB + 26.6% iodixanol). The sample is then loaded in a previously fabricated microfluidic channel.

Cell motility is visualized using the Nikon TI with a 10x (NA 0.3) objective. A stack of 10000 frames at 100 frames per second is acquired under phase contrast illumination with an Eosens 4cxp CMOS camera and StreamPix Multicamera software. The swimming speed of cells, the fraction of motile cells and the diffusion coefficient of non-motile cells (Supplementary Fig. S7) are measured using Differential Dynamic Microscopy^89^, an image analysis method based on Fourier image analysis, following the protocol described in Colin *et al*. ^29^.

To reconstruct the horizontal trajectories of the cells close to the surface, a stack of 2000 frames at 50 frames per second is acquired. Single 2D trajectories are reconstructed using the plugin ParticleTracking for ImageJ ^90^ (https://github.com/croelmiyn/ParticleTracking, DOI:10.5281/zenodo.7781880). Cell trajectories are categorized into runs and tumbles as previously described ^90^. To calculate the radius of curvature of the trajectories, only runs are considered. The parametric curvature *C*(*t*) is calculated for frame *t* as

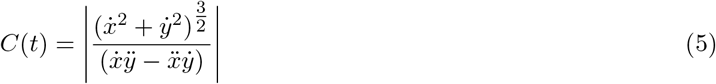

where the first and second derivatives of *x*(*t*) and *y*(*t*) are extracted from the cell tracking data by Taylor expansion. For this, a 50 frame sliding window is considered around time *t*, over which *x*(*t*^*′*^) and *y*(*t*^*′*^) are leastsquares fitted to a second-order polynomial to obtain their first and second derivatives, as given by Taylor’s expansion:

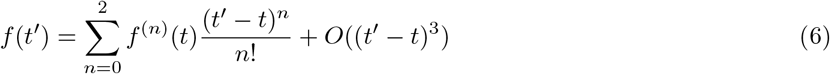

The mean curvature is computed for each run, and for each trajectory as the average of *C*(*t*) over all times *t* for which the cell is categorized as running. The radius is then the inverse of the curvature, *R* = 1*/ C*(*t*). When computing histograms, the mean radius *R* of a trajectory has a weight equal to the duration of this trajectory.

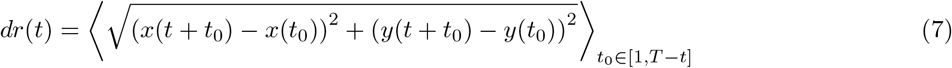

To estimate the circle persistence time of a trajectory of duration *T*, we compute the mean linear displacement and find the smallest lag time *t* = *τ* for which *dr*(*t*) exceeds the distance 3*R* with *R* the above defined radius of curvature, ie 1.5 times the average circle diameter. If *dr*(*t*) never exceeds 3*R*, we assign *τ* = *T*.

## Numerical simulations

All simulations were performed in a three dimensional rectangular box of width and length *W* and of height *H*. Boundary conditions are periodic on the lateral edges of the simulation box, and reflexive no-slip at the top and bottom walls. Cell volume fractions are *ϕ* = *nV/HW* ^2^, with *n* the number of cells and the cell volume *V* being twice higher for elongated cells. All parameter values are given in Table S2.

### Mean flow field generated by a circular swimmer

The motile cell is modeled as a pusher dipole of point forces, representing the cell body and the flagellum, both of magnitude *f*_0_*/*6*πη* = *a*^*S*^*v*_0_, with *η* the fluid viscosity, *a*^*S*^ the hydrodynamic radius of the cell body and *v*_0_ the swimming speed ^17^. The fluid flow **v**^*i←j*^ in position **x**_*i*_ generated by a point force **f**_*j*_ positioned in **x**_*j*_ is computed as

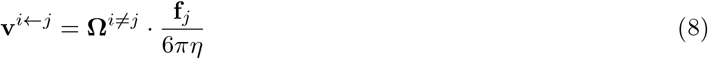

where **Ω**^*≠ j*^ is the normalized hydrodynamic-interaction tensor, i.e. the normalized Green function, in our geometry, of the incompressible Stokes equation:

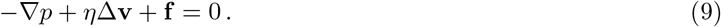

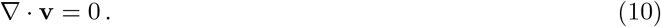

The tensor **Ω**^*≠ j*^ and therefore the hydrodynamic flows **v**^*i←j*^ are computed using the two-dimensional Fourier-space expansion of the hydrodynamic flow introduced in Mucha *et al*. ^71^, using the N log(N) algorithm of Hernandez-Ortiz *et al*. ^72^, which allows an efficient calculation of the fluid flow generated by multiple point forces in our geometry via the additivity of the solutions to Stokes equation, and which we implemented in Java (Supplementary notes). The dipole flow *v*^*D*^(**x**; **x**_0_, **u**) is the sum of the flow generated by the two point forces, +**u***f*_0_ located at **x**_0_ + *l*_*d*_**u** (cell body), and −*f*_0_**u** located at **x**_0_ − *l*_*d*_**u** (flagellum), with **x**_0_ the position of the center of the cell, **u** the unit vector pointing in the swimming direction and *l*_*d*_ the half dipole length. To describe circular surface swimmers, the dipoles are horizontal and located at a height *z*_*M*_ = 1 *µ*m from the surface over which they circulate. The mean flow field generated over one revolution is computed as the average of the dipole flow:

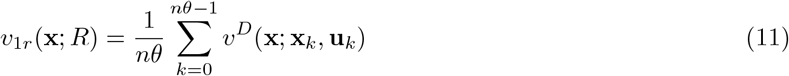

over *nθ* = 100 equally spaced positions of the dipole on a circle of radius *R*, **x**_*k*_ = (*R* cos *θ*_*k*_, *R* sin *θ*_*k*_, *z*_*M*_), with *θ*_*k*_ = 2*πk/nθ*, the dipole being tangential to the circle, **u**_*k*_ = (− sin *θ*_*k*_, cos *θ*_*k*_, 0). Because this configuration of forces is rotationally symmetric, the mean flow field only depends on the longitudinal (*z*) and radial (*r*) coordinates in the cylindrical coordinate system relative to the center of the circle, and has no azimuthal component.

The flow field was analyzed using MATLAB (2020a). We define the inward flux generated by the circular swimmer at a distance *r* by:

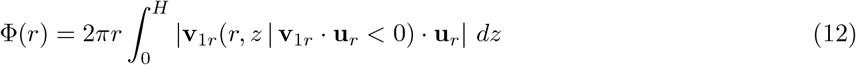

with **u**_*r*_ the radial unit vector. The recirculation radius is defined as the distance *r* which maximizes Φ, corresponding to the position of the center of the recirculation whirl. The recirculation flux is defined as the maximal flux max_*r*_ Φ(*r*).

The vertically averaged *v*_1r_ (**x**; *R*) of Fig. 4d,e is computed as 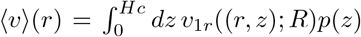; *R*)*p*(*z*), with the fitted tracer distribution 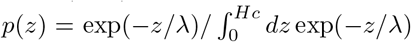, *λ* = 10 *µ*m the experimental decay length and *Hc* = 20 *µ*m the height of the confocal volume.

### Distribution of tracers in the mean flow field of a population of circular swimmers

The mean flow field created by a population of *n*_*Mc*_ (250 to 5 10^4^) circular swimmers sums the mean flows *v*_1*r*_ of the individual circular swimmers. Swimmers are assigned with equal probability to the top (*z* = *H* − *z*_*M*_) or bottom (*z* = *z*_*M*_) wall, mimicking experiments (Supplementary Fig. S7b). The centers of the circles are distributed uniformly in the horizontal (*x, y*) directions. The circle radius is drawn from a double log normal distribution *p*(*R*), which was parametrized to fit the experimental distribution of circle radii for normal-sized and elongated cells (Supplementary Fig. S11a). We perform a simplified Brownian dynamics simulation of the motion of *n*_*NM*_ (5 10^3^ to 2 10^4^) non-motile cells that are subjected to the mean flow field of the circles, gravity and Brownian motion. The evolution of the position of tracer *i* obeys

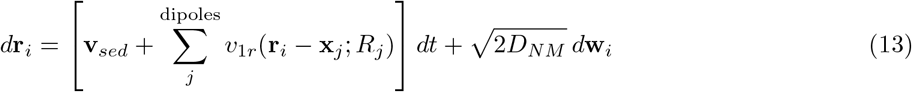

with **v**_*sed*_ the downward-oriented sedimentation speed, **x**_*j*_ and *R*_*j*_ the center and radius of circle *j*, which are kept fixed throughout the <u>sim</u>ulation, *dt* the simulation time step, *D*_*NM*_ the diffusion coefficient of the non-motile cells, and 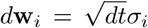 the normalized Brownian noise. The Gaussian-distributed three-dimensional random variable *σ*_*i*_ has unit variance and no cross-correlation with *σ*_*j≠ >i*_. The initial distribution of non-motile cells is uniform horizontally and obeys the Boltzmann distribution in confined geometry along the vertical axis

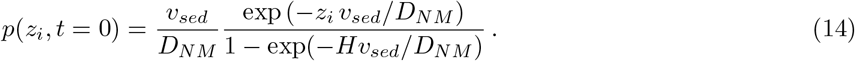

Steric interactions, direct hydrodynamic interactions between non-swimmers and the effect of confinement on self-diffusion are neglected (Supplementary note). The equation of motion is time integrated using an Euler scheme, with *dt* = 0.1 s in equivalent real time. The configuration of the system is saved every step, for a total duration of the simulation of 1000 steps. Because the circle configuration and thus the velocity field are fixed, the time elapsed since the beginning of the simulation can be interpreted as a persistence time of the circles, and the configuration after this time as a realization of the structure of the population of non-motile cells in the mixture for said persistence time.

A pseudo image of the suspension is produced for each time step of the simulation from the positions of the non-motile tracers. It renders the microscopy image that is expected under our experimental conditions for the given non-motile cells configuration, assuming a Gaussian point spread function for our objective. Each particle, positioned in (**r**_**i**_ = (*x*_*i*_, *y*_*i*_), *z*_*i*_) contribute to the intensity in position **r** = (*x, y*) in the rendered image at *h* = 8 *µ*m above the bottom of the channel as:

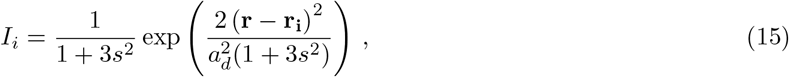

with *s* = (*z*_*i*_ − *h*)*/dh* the distance to the focal plane, normalized to the depth of field *dh*. The images are printed with a pixel size of px_*sim*_ = 1 *µ*m, which is larger than in our experimental setup, to save computer memory. They are then analyzed with the same algorithms as the experimental data. We used a box size of 32 × 32 *µ*m^2^ to compute the heterogeneity level as *σ*^2^ */Nσ*^2^, averaged over the persistence time window [500700] steps, and normalized to its value in the purely passive homogeneous control similarly to experiments. The value of 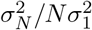 in the simulated passive system is 8.9, smaller than in experiments owing to the smaller pixel size there (1 px_*exp*_ = 0.325 *µ*m).

The simulations for each parameter set are repeated six times and quantifications are averaged over these repeats. This relatively small number of repeats is sufficient because the heterogeneity level varies little between independent repeats. The fully parallelized java code for running the simulations is available on Github (https://github.com/croelmiyn/SimulationMixtures, DOI:10.5281/zenodo.7919119).

## Acknowledgements

The authors acknowledge support from the Deutsche Forschungsgemeinschaft (Grant No. CO 1813/2-1). The authors thank L. Laganenka, O. Schauer and G. Scarinci for providing plasmids, G. Malengo for technical microscopy support, Chia-Fu Chou for helping with wafer fabrication, and V. Sourjik, S. Murray and H. Niederholtmeyer for helpful discussions.

## Data Availability

The data that support the findings of this study are available from the corresponding author upon reasonable request.

## Code Availability

Simulation code is available on Github at: https://github.com/croelmiyn/SimulationMixtures, DOI:10.5281/zenodo.7919119

Previously published codes for Differential Dynamic Microscopy and Image velocimetry as well as particle tracking are available on Github at:

https://github.com/croelmiyn/FourierImageAnalysis, DOI:10.5281/zenodo.3516258 https://github.com/croelmiyn/ParticleTracking, DOI:10.5281/zenodo.7781880

## Supplementary Information to

### Supplementary Figures

**Figure S1:**
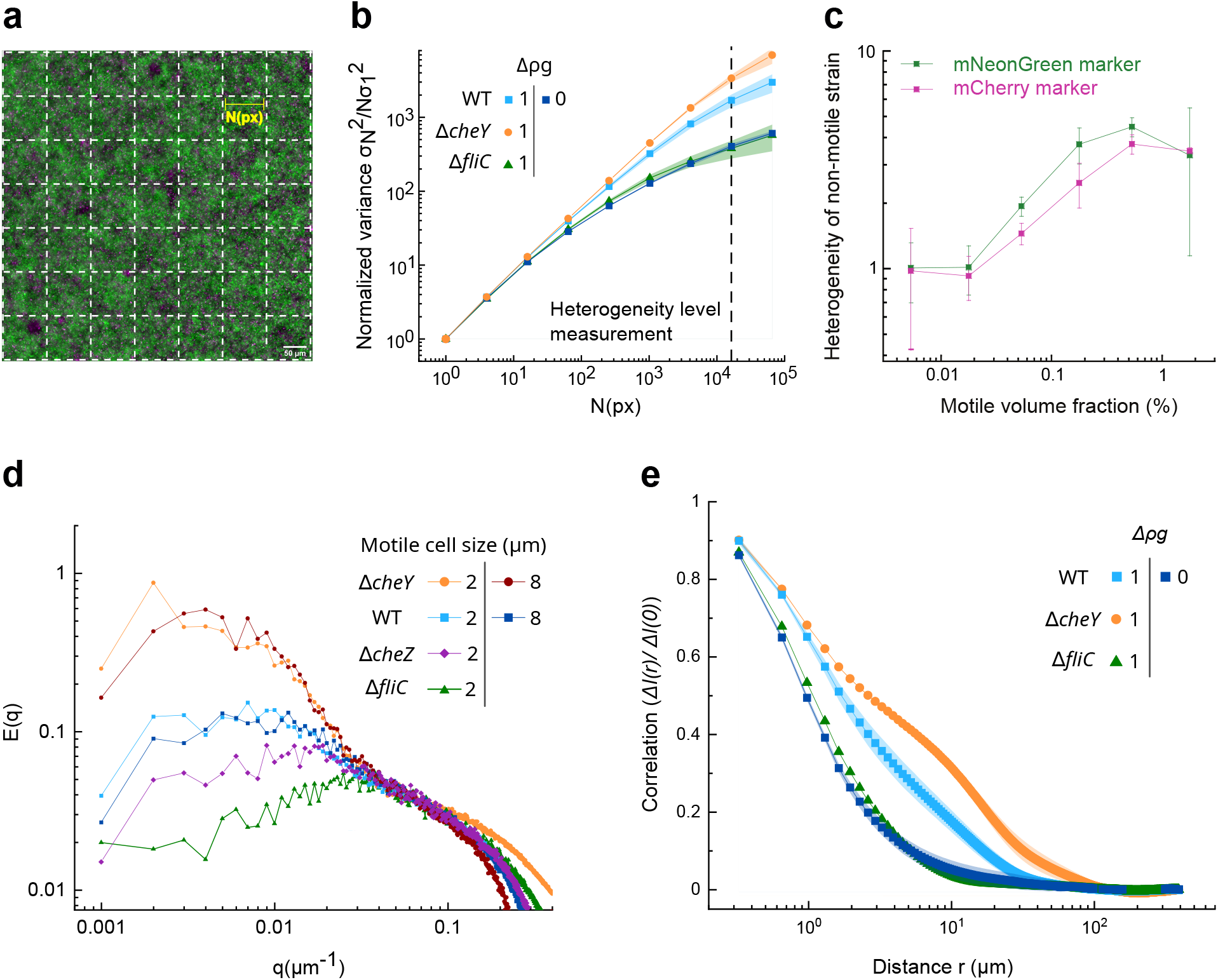
Characterization of the density fluctuations. a) Fluorescent image divided into nonoverlapping squares of area *N* px^2^. b) Normalized variance of the fluorescence intensity of the non-motile strain 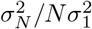 as a function of the number *N* of pixels of the division square, in the binary mixture *ϕ*_*NM*_ = 5.3%, *ϕ*_*M*_ = 0.18%. The dotted line represents the value of *N* for which heterogeneity level is calculated. c) Comparison of the heterogeneity level as a function of the density of motile cells, for a fixed non-motile cell density *ϕ*_*NM*_ = 5.34%, obtained when the non-motile cells are marked with either the mNeonGreen or the mCherry marker as indicated, the motile cells being marked with the other one. d) Structure factor E(q) of the non-motile strain in the binary mixture *ϕ*_*NM*_ = 5.3%, *ϕ*_*M*_ = 0.18% with the different strains used in this study. e) Spatial autocorrelation function of the fluorescence intensity for the non-motile cells in the binary mixture *ϕ*_*NM*_ = 5.3%, *ϕ*_*M*_ = 0.18%, featuring a short scale decay around 1 *µ*m, corresponding to the cell size, and a second non-exponential decay at ∼ 10-50 *µ*m length scales in heterogeneous samples. b-e) The data points consist of the geometric mean of three biological replicates and error areas or bars represent the standard deviation.

**Figure S2:**
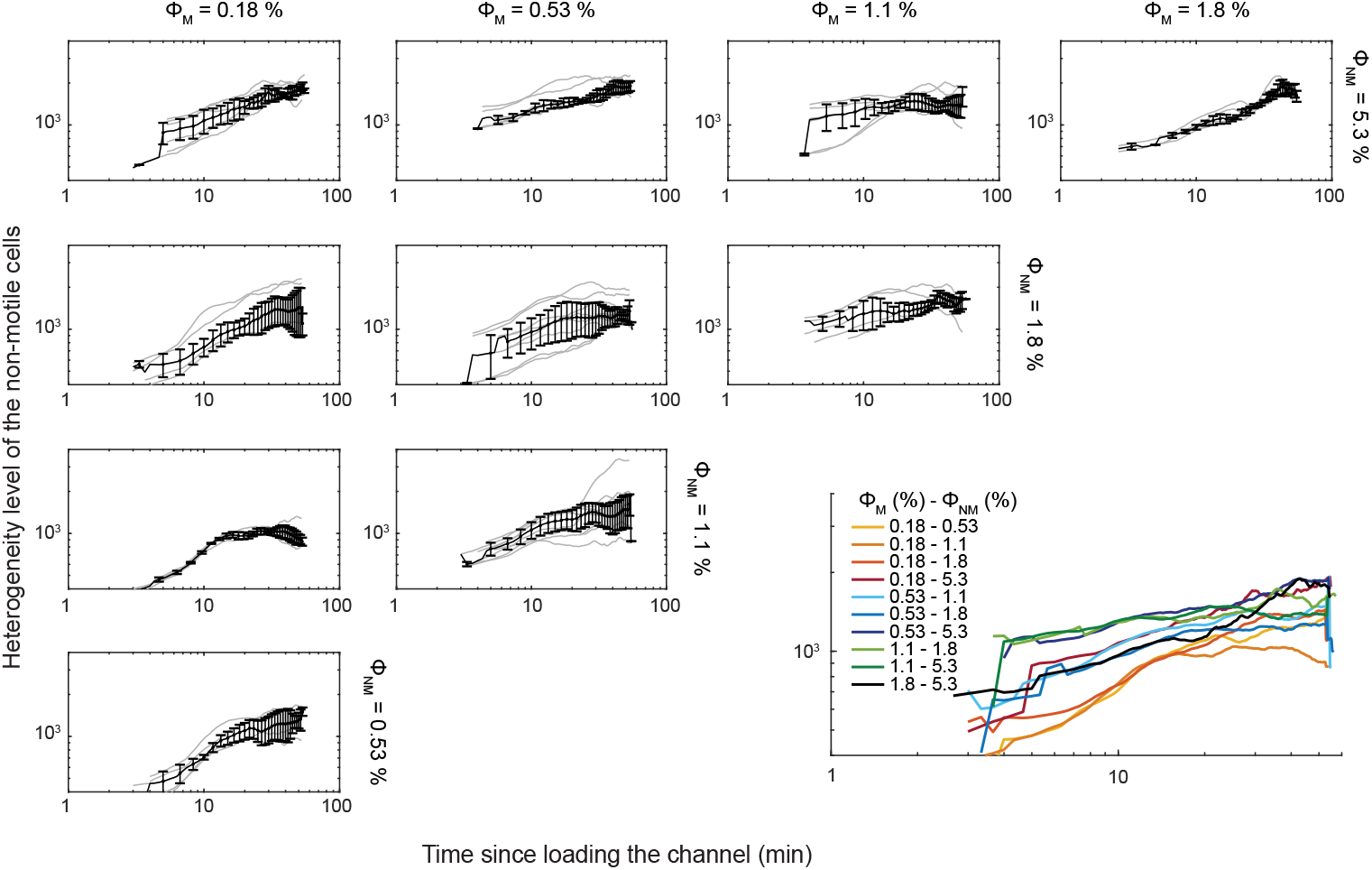
Dynamics of emergence of the density fluctuations. The heterogeneity level of the nonmotile strain of the mixture is shown as a function of the time since the mixture was loaded in the channel of the microfluidic chip, for different combinations of volume fractions of motile and non-motile cells. Displayed is the geometric mean of three independent replicates and error bars represent the standard deviation.

**Figure S3:**
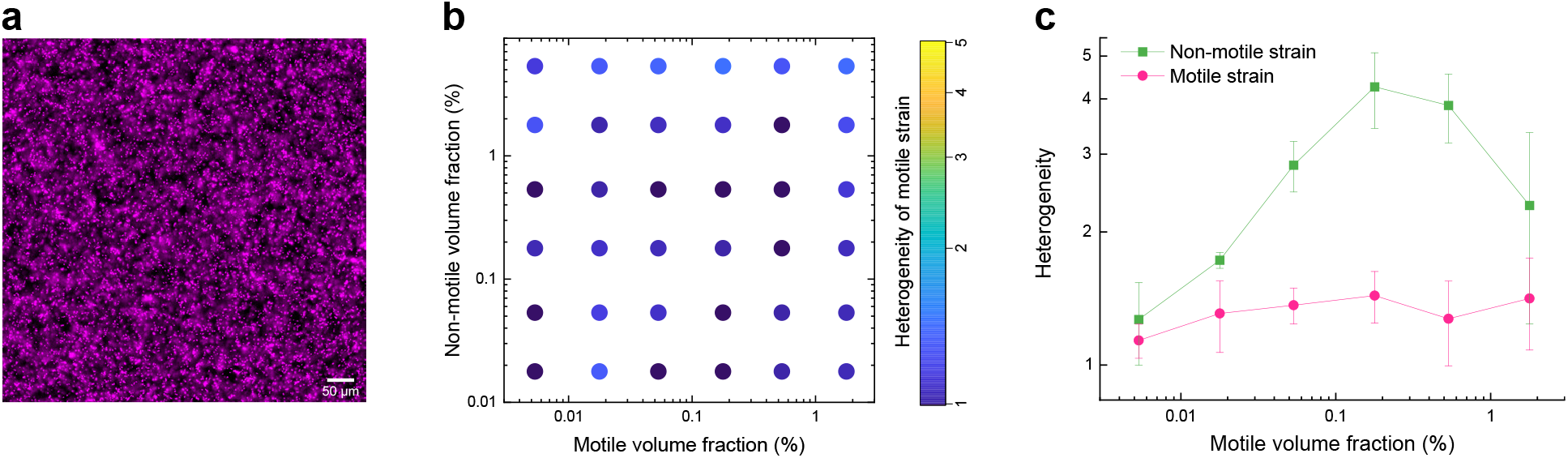
Behavior of the motile strain in the binary mixture. a) Microscopy image of the motile strain (WT *E*.*coli* mCherry) the binary mixture *ϕ*_*NM*_ = 5.3%, *ϕ*_*M*_ = 0.18%. b) Phase diagram of the density fluctuations of the motile strain in binary mixtures of bacterial populations as a function of the volume fraction (in %) of each strain. The heterogeneity level, which quantifies density fluctuations, is represented by a color scale. Each point represents the geometric mean of three independent replicates. c) Comparison between the heterogeneity level of the motile and the non-motile strains as a function of the volume fraction of the motile strain. The volume fraction of the non-motile strain is kept constant at 5.34%. Each point represents the geometric mean of three independent replicates and error bars indicate the standard deviation.

**Figure S4:**
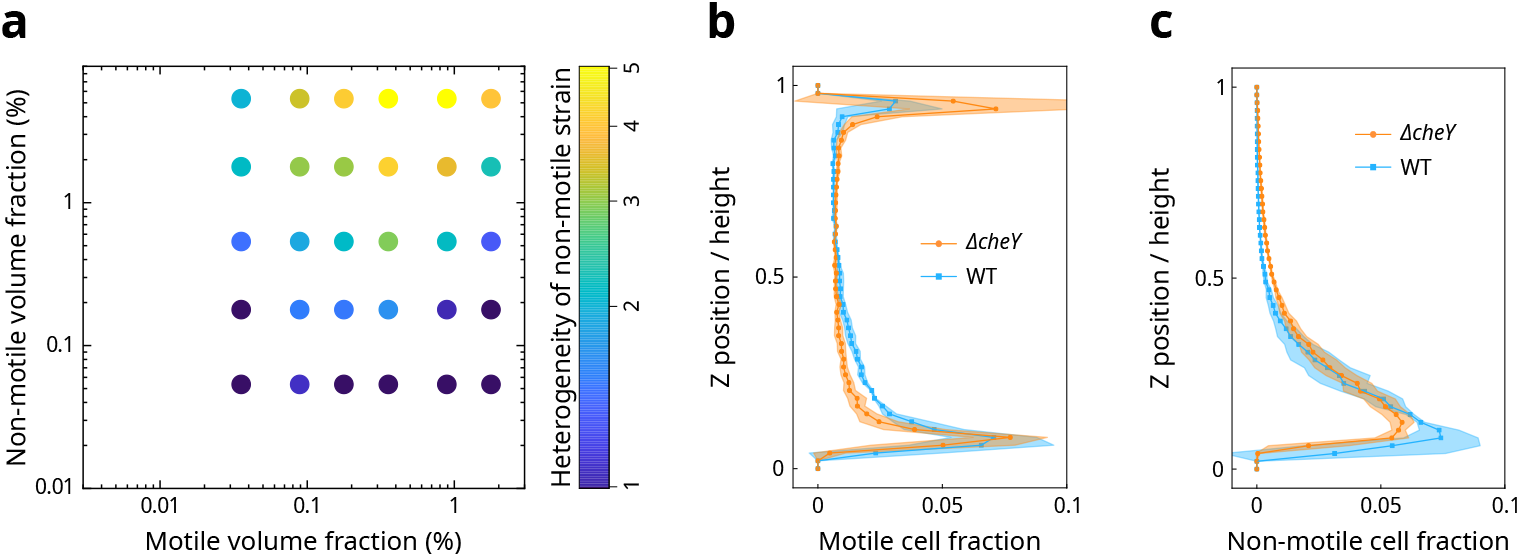
Effect of chemotaxis and tumbling. a) Phase diagram of the density fluctuations of the nonmotile strain in binary mixtures with *cheY* ** as the motile strain. The quantification of the density fluctuations as heterogeneity level is represented by a color scale. Each point represents the mean of three replicates. b-c) Motile (b) and Non-motile (c) cell fraction (abscissa), measured by cell counting in confocal microscopy, as a function of the vertical position in a channel of 50 *µ*m of height (ordinate), in the binary mixtures with *ϕ*_*NM*_ = 5.3% and *ϕ*_*M*_ = 0.18%, with the Δ*cheY* and WT motile strains. Each point represents the mean and error area the standard deviation over three replicates.

**Figure S5:**
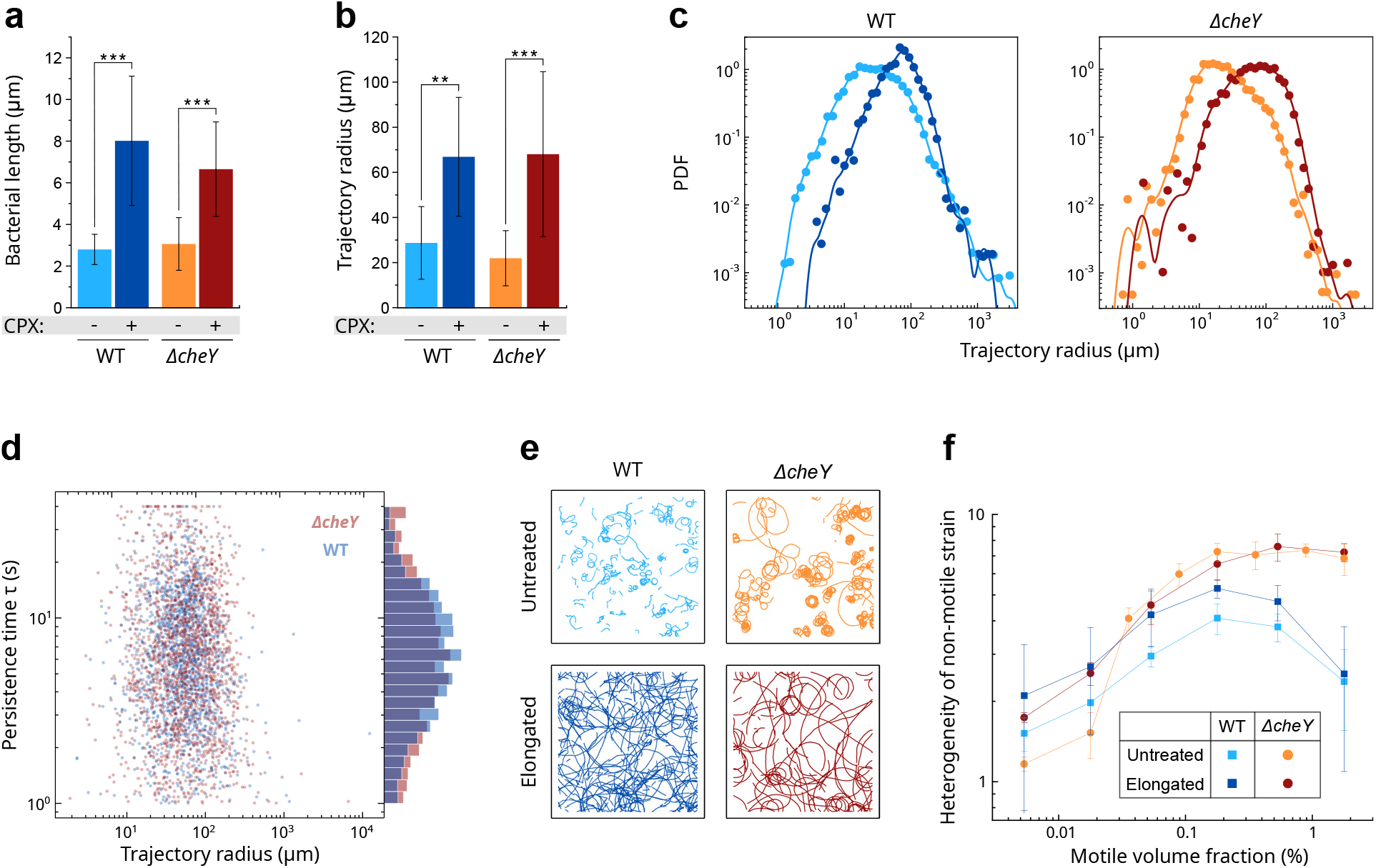
Effect of cell elongation. a-b) Median (a) bacterial cell length and (b) trajectory radius described by the motile cells (WT: run and tumble, Δ*cheY* : non-tumbling), untreated and elongated, swimming close to the surface. The error bars represent the median absolute deviation. Welch’s t-test: \*\**p <* 0.001, \*\*\**p <* 0.0001. c) Probability distribution of the radii from the circular trajectories. The radii are weighted by the trajectory length. a-c) The cell length and the trajectory radius were measured as described in the Methods section. Displayed are 1828 (WT CPX+), 1799 (Δ*cheY* CPX-), 4475 (WT CPX-) and 4137 (Δ*cheY* CPX-) individual swimming trajectories from 4 (CPX+ data) or 8 (CPX- data) biological replicates. d) Scatter plot of the persistence time *τ* of individual circular swimming trajectories of elongated WT and Δ*cheY* motile cells as a function of the radius of curvature of the trajectory, and probability density distribution of *τ* (right side). e) Swimming trajectories of untreated and elongated cells, both the run and tumble strain (WT) and the non-tumbling strain (Δ*cheY*). f) Comparison between the heterogeneity level of the elongated and untreated cells in both strains (WT and Δ*cheY*). The volume fraction of the non-motile strain is kept fixed to 5.34% and the volume fraction of the motile strain is systematically varied. Each point represents the geometric mean of three independent replicates and error bars indicate the standard deviation.

**Figure S6:**
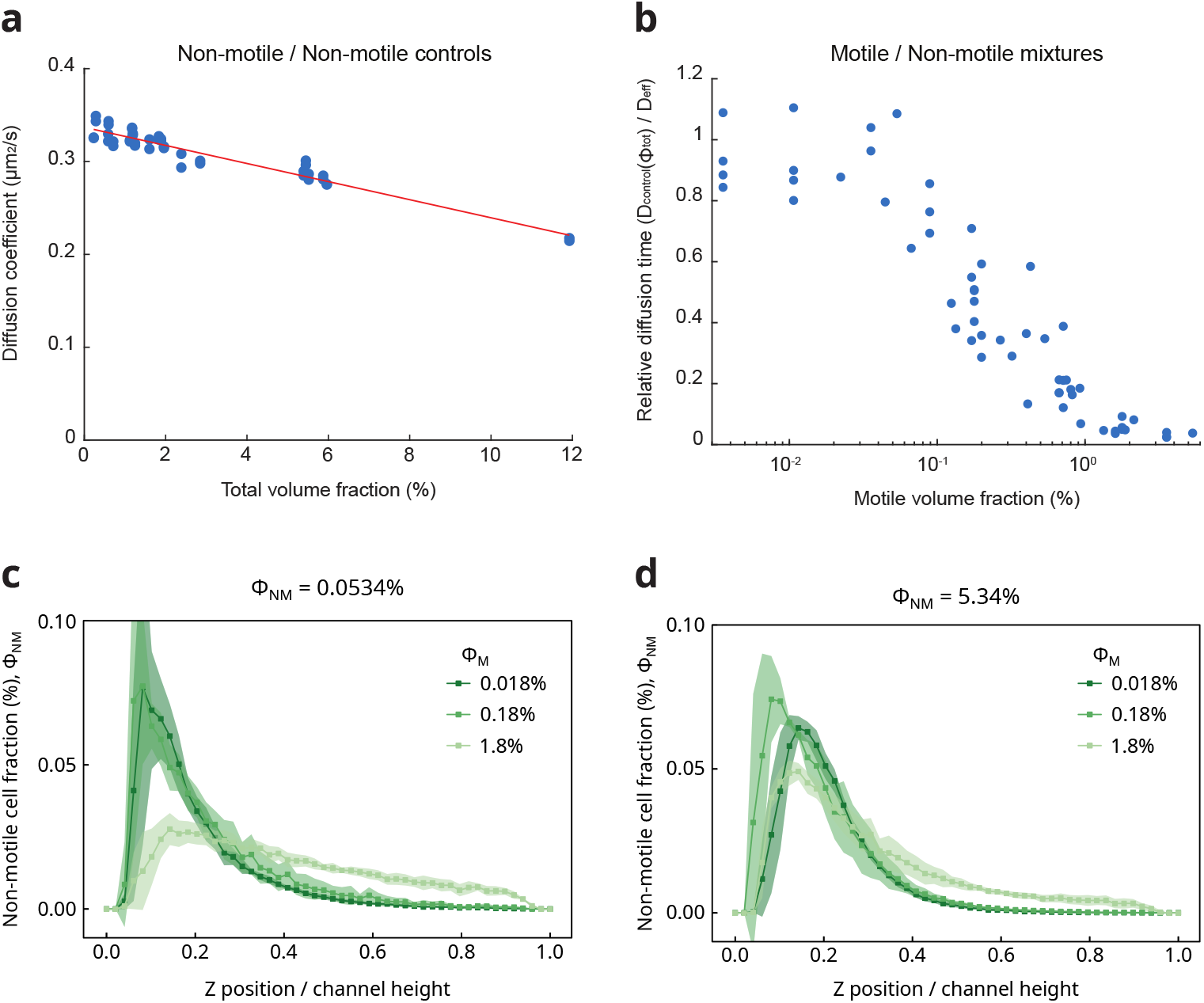
Effect of the motile cells on the diffusion of the non-motile cells. a-b) Diffusion of the non-motile cells measured using Differential Dynamic Microscopy. a) Diffusion coefficient of the non-motile cells in absence of motile cells (control) as a function of the total volume fraction of cells in the population. The straight line is a linear fit. b) Inverse of the diffusion coefficient of the non-motile cells (normalized by its value in the non-motile only case at the same density) as a function of the volume fraction of run and tumble (WT) motile cells in the mixtures. The increase in the diffusion of the non-motile cells becomes noticeable for volume fractions of motile cells above 0.03%. c-d) Non-motile cell fraction (abscissa), measured by cell counting in confocal microscopy, as a function of the vertical position in a channel of 50 *µ*m of height (ordinate), for varying volume fractions of the motile strain (WT) in the mixture, in the cases of a low (c) or a high (b) total volume fraction of non-motile cells.

**Figure S7:**
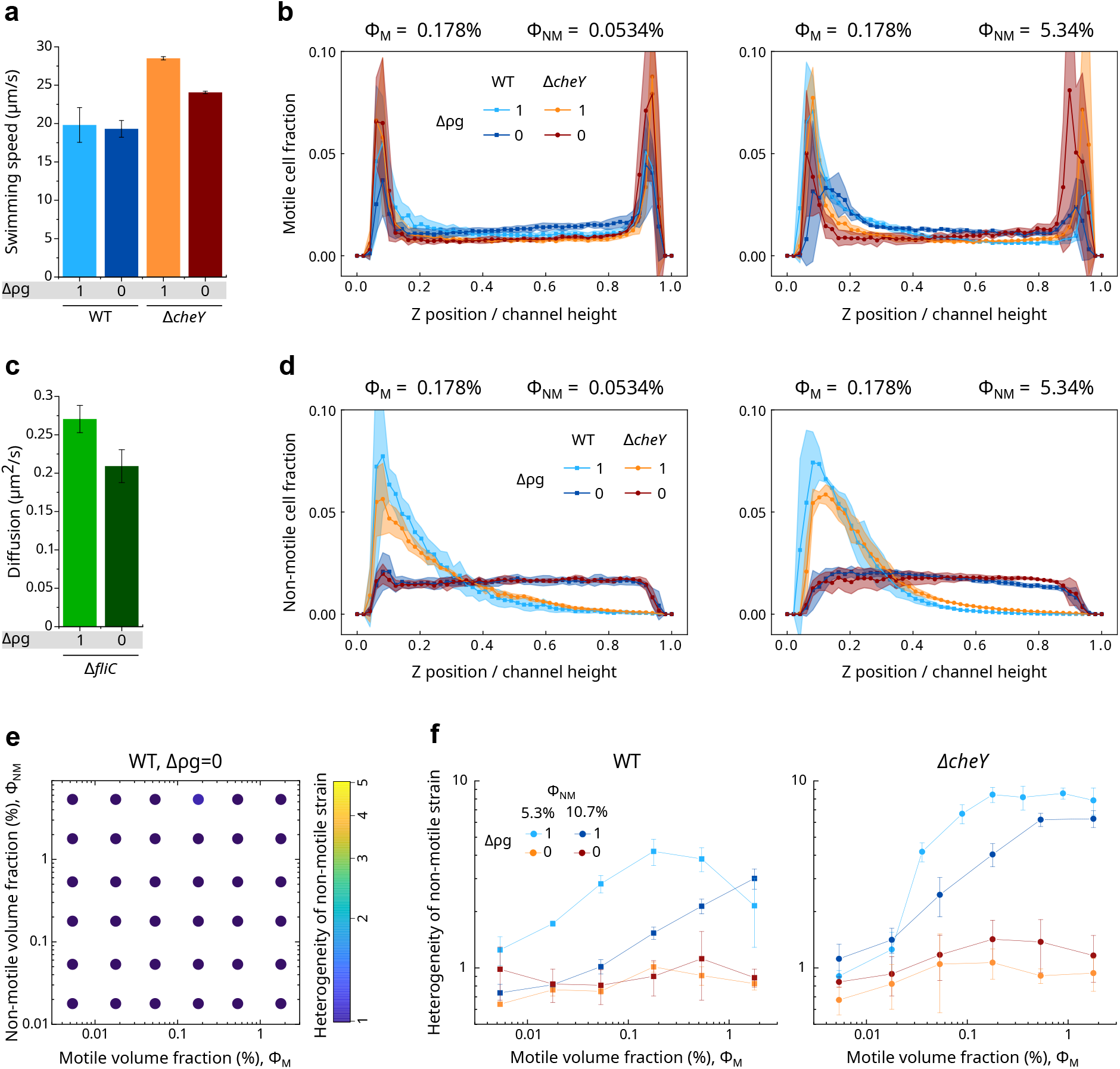
Effect of Iodixanol and neutral buoyancy. (a-d) Effect of Iodixanol on cell motion and position. (a) Swimming speed of motile cells (WT and Δ*cheY*) and (c) diffusion coefficient of non-motile cells alone (Δ*fliC*) were measured using Differential Dynamic Microscopy in regular motility buffer (Δ*ρg* = 1) and in neutrally buoyant medium, i.e. motility buffer with 26.6% of Iodixanol (Δ*ρg* = 0). (b, d) Vertical localization of the (b) motile and (d) non-motile cells in the binary mixture *ϕ*_*NM*_ = 5.3%, *ϕ*_*M*_ = 0.18% and in a channel of 50 *µ*m of height. The type of motile cells (WT or Δ*cheY*) and the suspending medium conditions (Δ*ρg* = 1 or 0) are as indicated. The cell count, measured from confocal microscopy Z-stacks, was normalized to total cell number to obtain the cell fraction, shown as a function of the Z position. e) Phase diagram of the density fluctuations of the non-motile strain (volume fraction in ordinate) in mixture with the WT motile strain (volume fraction in abscissa) in density-matched suspending media (Δ*ρg* = 0). The heterogeneity level, which quantifies density fluctuations, is represented by a color scale. f) Effect of increased non-motile volume fraction (*ϕ*_*NM*_) on the density fluctuations. The heterogeneity of the non-motile strain is plotted against the volume fraction of the motile strain for two *ϕ*_*NM*_ ; 5.3 % and 10.7 %, for two sedimentation regimes; Δ*ρg* = 1 and Δ*ρg* = 0, and for two motile strains; WT and Δ*cheY*. a-f) Each point represents the mean (a,c), binned mean (b,d) or geometric mean (e-f) of three independent replicates and error bars or areas indicate the standard deviation.

**Figure S8:**
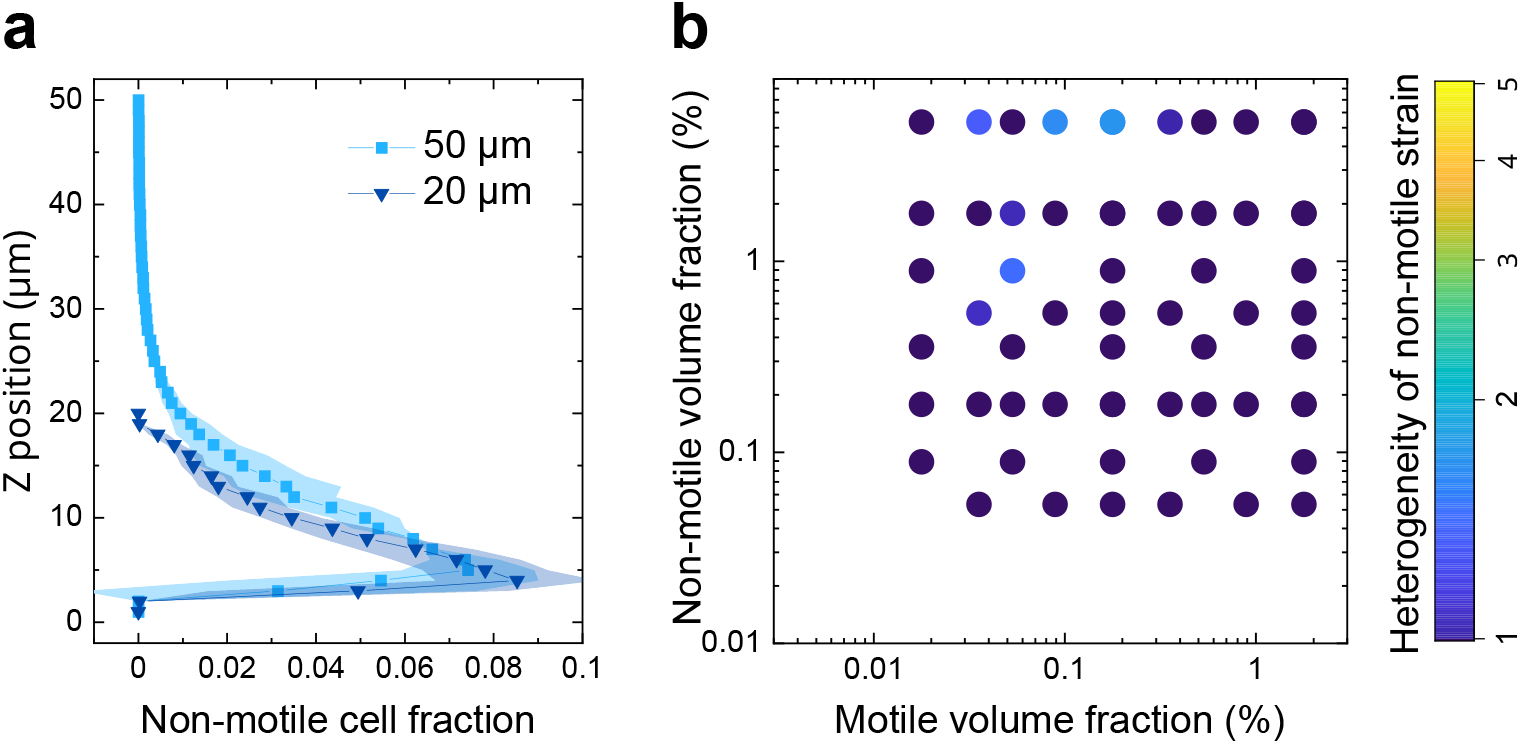
Effect of vertical confinement in a 20 *µ*m high channel. a) Vertical distribution of nonmotile cells in the binary mixture *ϕ*_*NM*_ = 5.3 %, *ϕ*_*M*_ = 0.18 %, with a WT motile strain, for two channel heights (*h* = 50 *µm* and *h* = 20 *µm*). b) Phase diagram of the density fluctuations of the non-motile strain in binary mixtures with WT as the motile strain in channels of 20 *µm* in height. The quantification of the density fluctuations as heterogeneity level is represented by a color scale. Each point represents the mean of three replicates.

**Figure S9:**
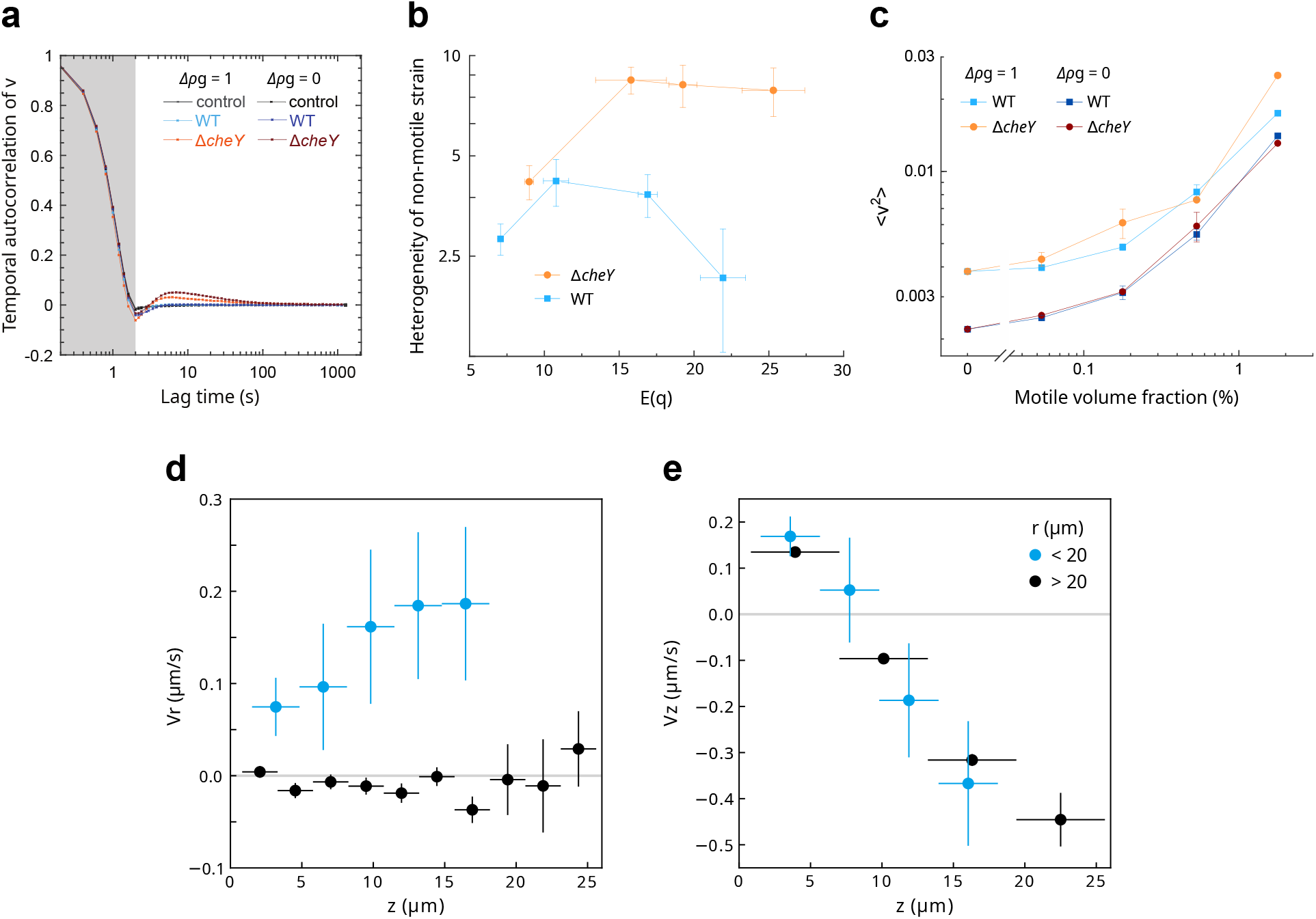
Dynamics of the non-motile cells in the density fluctuations. a) Temporal autocorrelation of the non-motile cell velocity in the binary mixture *ϕ*_*NM*_ = 5.3%, *ϕ*_*M*_ = 0.53% with the indicated motile cells (WT, Δ*cheY* and control (Δ*fliC*)) and buoyancy (Δ*ρg* = 1 and Δ*ρg* = 0). The grey area indicates the width of the sliding window that was used to estimate the velocity. The non-motile cells’ velocity shows long-time (*t >* 5 s) positive correlations only in presence of Δ*cheY* motile cells. b) The heterogeneity level of the nonmotile strain in mixture with two different motile strains, WT and Δ*cheY*, as a function of the peak of the flow structure factor E(q) (Fig. 4b) that quantifies the amplitude of the flow patterns. The volume fraction of the non-motile strain is kept to *ϕ*_*NM*_ = 5.3% and the one of the motile strain increases with E(q) (Fig. 4b). Each point represents the mean over a triplicate of the peak of E(q) with the standard deviation as the error bars, and the geometric mean of the heterogeneity level with the standard deviation as the error bars. c) Kinetic energy ⟨*v*^2^⟩ of the non-motile strain as a function of the motile volume fraction *ϕ*_*M*_ for mixtures with different motile strains (WT and Δ*cheY*) and buoyancies (Δ*ρg* = 1 and Δ*ρg* = 0). d) Vertical (*V*_*z*_) and e) radial (*V*_*r*_) velocity relative to the center of the depletion zones (hole) of the non-motile tracer cells as a function of their vertical position (*z*), for cells that are less (blue), or more (black), than 20 *µ*m away from the hole center (*r*). Each data point represents the binned mean, the vertical error bars represent the standard error, and the horizontal error bars represent the bins over which the mean was calculated. The data comprises 1591 individual 3D trajectories.

**Figure S10:**
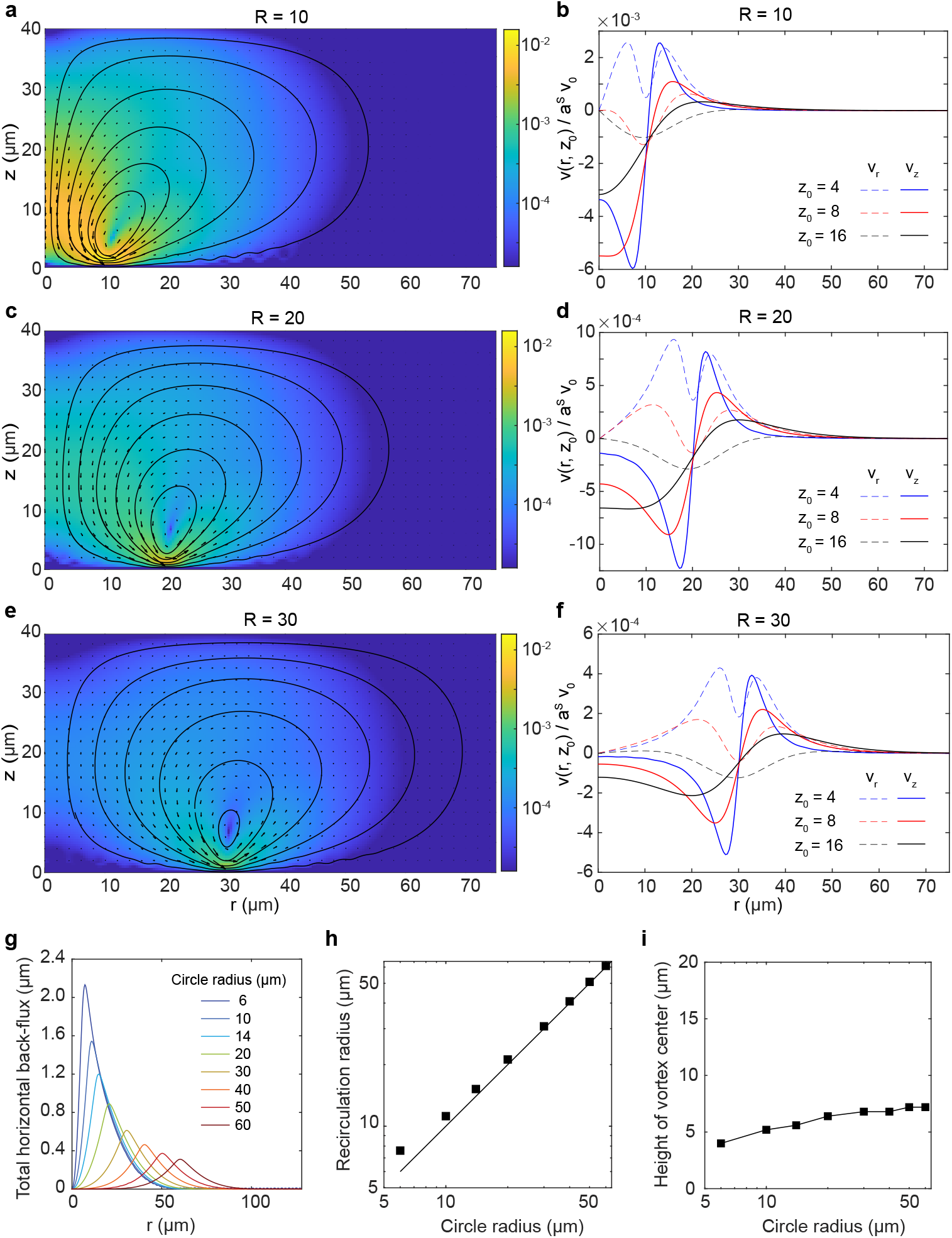
Computed flow field created by a circular swimming dipole for several circle radii *R*, situated *z*_*M*_ = 1 *µ*m above the bottom in a channel of height *H* = 40 *µ*m. a-f, Shown are the flow fields and some streamlines as a function of the horizontal distance to the center of the circle *r* and the height within the channel *z* (a,c,e), and the normalized vertical(*v*_*z*_) and radial (*v*_*r*_) velocities at the indicated heights *z*_0_ (b,d,f), for the indicated radii. The normalized velocity *v*(*r, z*)*/a*^*S*^*v*_0_ is in *µ*m^*−*1^. g, Total back-flux Φ(*r*)*/a*^*S*^*v*_0_ (equation 12) through a cylinder of radius *r* centered on the circle of indicated radius (R). The back-flux is maximum for *r* corresponding to the position of the eye of the recirculation vortex, called the recirculation radius. The maximum is the recirculation flux plotted in Fig. 5c. h, Recirculation radius and i, height of the eye of the recirculation vortex, as a function of the radius of the circle.

**Figure S11:**
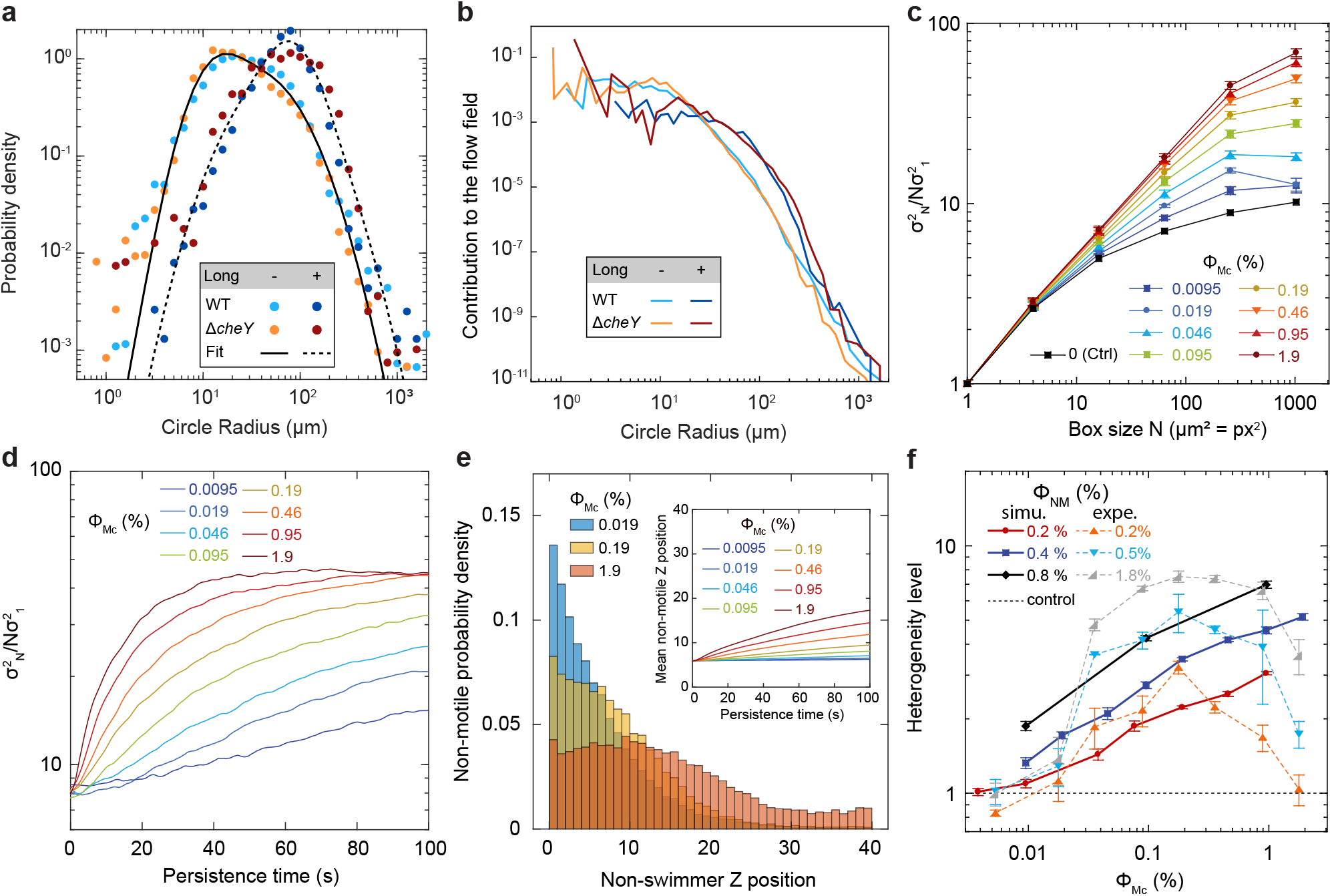
Supplementary data to the simulations of mixtures of motile circular swimmers (Mc) and non-motile tracer cells (NM). a) Comparison between the ansatz probability density of a circle having a given radius (Fit) and the experimental distributions for normal (Long -) and elongated cells (Long +) for indicated cells. b) Estimation of the contributions to global flows of a cell swimming with a given radius of curvature. The estimation is computed as *l*_*p*_*a*^*S*^*v*_0_*p*(*R*)*/R*^3^, i.e. the product of the dipole strength *l*_*p*_*a*^*S*^*v*_0_, with the dipole length *l*_*p*_, the cell body hydrodynamic radius *a*^*S*^ and the swimming speed *v*_0_, times the probability *p*(*R*) of an observing radius *R* times the typical magnitude of the fluid velocity produced by the circular swimmer for this radius (⟨|*v*_1*r*_|⟩ ∝ 1*/R*^3^). c) Heterogeneity measure 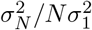 (equation 2) of the NM cells (volume fraction *ϕ*_*NM*_ = 0.4 %) as a function of the box size *N* for the indicated volume fractions of motile circular swimmers of normal size (*ϕ*_*Mc*_), averaged over the persistence time range *τ* = [50, 70] s. d) Heterogeneity measure 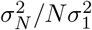 as a function of the time since the start, from a homogeneous state, of the simulation (persistence time) for indicated *ϕ*_*Mc*_ for *N* = 256. The heterogeneity level is the average of 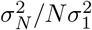 over *τ* = [50, 70] s for *N* = 256. e) Vertical distribution of non-motile cells for the indicated volume fractions of motile cells *ϕ*_*Mc*_ after a persistance time *τ* = 60 s. (inset) Average vertical position of the NM cells (*ϕ*_*NM*_ = 0.4 %) throughout the simulation for indicated *ϕ*_*Mc*_. The increase for larg *ϕ*_*Mc*_ reflect the increased effective diffusion. f) Heterogeneity level as a function of the volume fraction of motile circular swimmers *ϕ*_*Mc*_, for different values of the volume fraction of non-motile cells (*ϕ*_*NM*_), compared to experiments with the non-tumbling mutant (expe.). Control is a homogeneous distribution of cells. c-f) For simulation data, points are the mean and error bars (c,f) the standard error of the mean over 6 independent repeats. f) Experimental data are the geometric mean and error bars the standard deviation over 3 biological replicates.

## Supplementary Tables

**Table S1:**
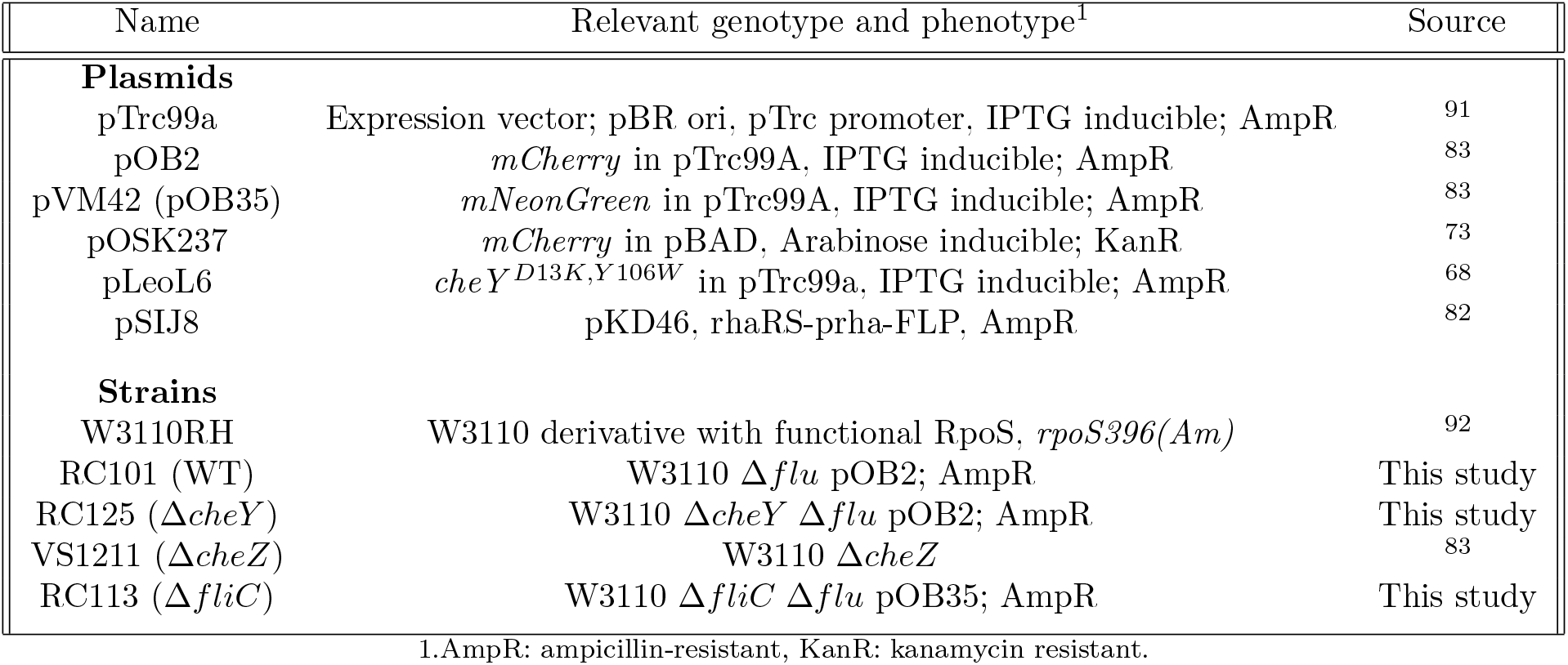
Plasmids and strains used in this study.

**Table S2:**
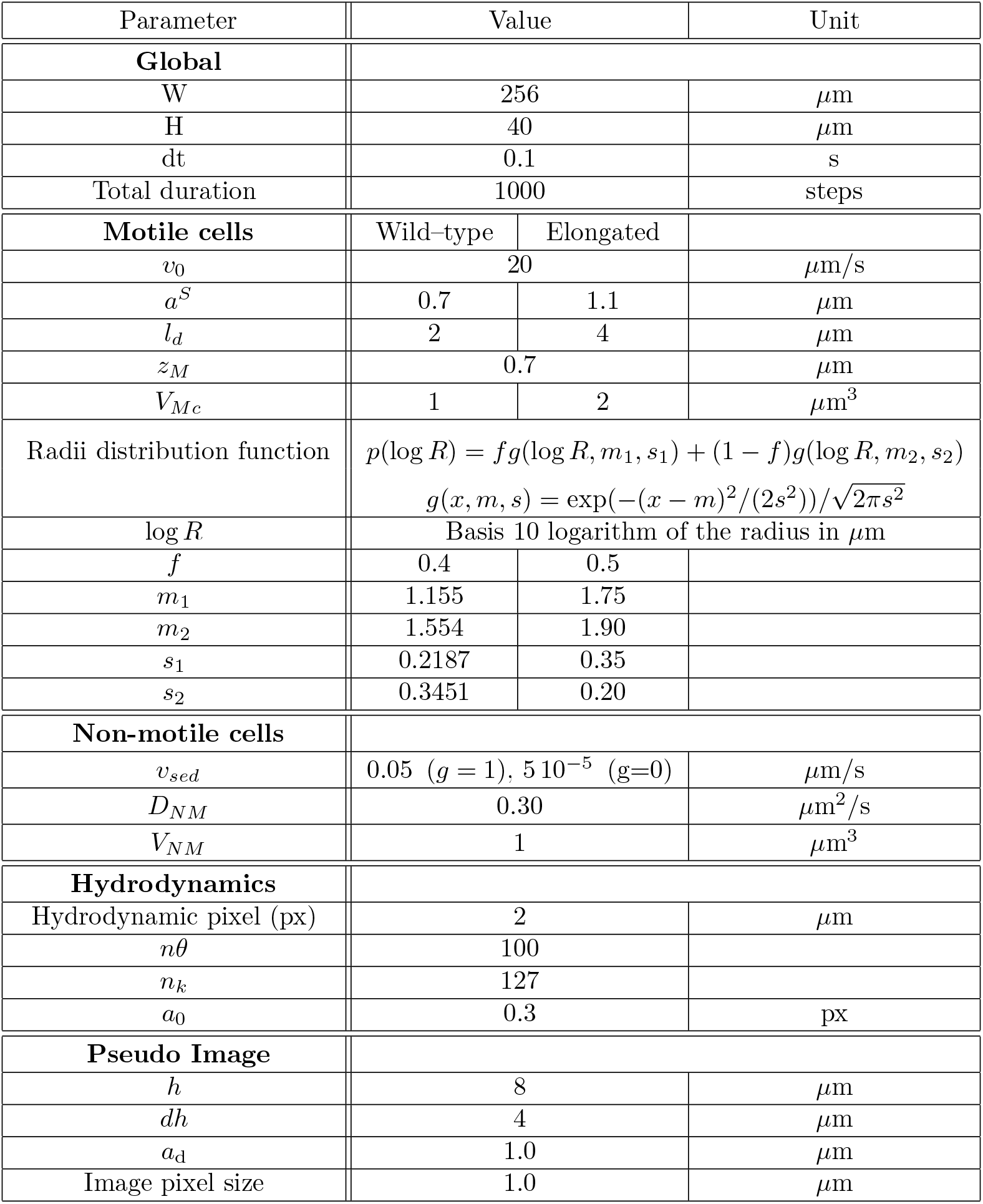
Coefficient values used in the numerical simulation

**Table S3:** Expressions for the *ca*^*b,n*^(*l, m*) coefficients, with the constant *C*_*h*_ = 3*π* exp(*kh*)*/*(*kW* ^2^(2*e*^4*hk*^(1 + 8*h*^2^*k*^2^) − *e*^8*hk*^ − 1)) and **k** = (*l, m*), *k* = |**k**| being non-zero.

| i,j/n | 0 | 1 | 2 | 3 | 4 |
| --- | --- | --- | --- | --- | --- |
|  | **b=1** |  |  |  |  |
| 1,1 | $C_h/(k^2(1-e^{4hk}))$ | $kl^2(1+e^{8hk}(1-4kh)+e^{4hk}(-2+4kh))$ | $(hkl^2+k^2+m^2)+e^{4hk}(2kl^2h-4h^2k^2(l^2+8m^2)-2(k^2+m^2))+e^{8hk}(-3l^2kh+4h^2k^2l^2+k^2+m^2)$ | $kl^2(e^{10hk}+e^{6hk}(-2-4kh)+e^{2hk}(1+4kh))$ | $e^{10hk}(hkl^2-k^2-m^2)+e^{6hk}(2kl^2h+4h^2k^2(l^2+8m^2)+2(k^2+m^2))+e^{2hk}(-3hkl^2-4h^2k^2l^2-k^2-m^2)$ |
| 1,2 | $C_h ml/(k^2(e^{4hk}-1))$ | $-k+e^{8hk}(-k+4hk^2)+e^{4hk}(2k-4hk^2)$ | $(-kh+1)+e^{8hk}(3hk-4h^2k^2+1)+e^{4hk}(-2kh-28h^2k^2-2)$ | $e^{10hk}(-k)+e^{2hk}(-k-4hk^2)+e^{6hk}(2k+4hk^2)$ | $e^{10hk}(-1-hk)+e^{2hk}(3hk+4h^2k^2-1)+e^{6hk}(-2hk+28h^2k^2+2)$ |
| 1,3 | $C_h l$ | $1+e^{4hk}(-1-4hk)$ | $h+e^{4hk}(-h+4h^2k)$ | $e^{6hk}(1)+e^{2hk}(-1+4kh)$ | $e^{6hk}(h)+e^{2hk}(-h-4h^2k)$ |
| 3,1 | $C_h l$ | $1+e^{4hk}(-1+4hk)$ | $h+e^{4hk}(-h-4h^2k)$ | $e^{6hk}(1)+e^{2hk}(-1-4kh)$ | $e^{6hk}(h)+e^{2hk}(-h+4h^2k)$ |
| 3,2 | $C_h m$ | $1+e^{4hk}(-1+4hk)$ | $h+e^{4hk}(-h-4h^2k)$ | $e^{6hk}(1)+e^{2hk}(-1-4kh)$ | $e^{6hk}(h)+e^{2hk}(-h+4h^2k)$ |
| 3,3 | $C_h$ | $(-k)+e^{4hk}(k+4hk^2)$ | $(1-kh)+e^{4hk}(-3hk-4h^2k^2-1)$ | $e^{6hk}(k)+e^{2hk}(-k+4k^2h)$ | $e^{6hk}(hk+1)+e^{2hk}(3hk-4h^2k^2-1)$ |
|  | **b=0** |  |  |  |  |
| 1,1 | $C_h/(k^2(1-e^{4hk}))$ | $-kl^2e^{10hk}+e^{6hk}(2kl^2+4k^2l^2h)+e^{2hk}(-l^2k-4l^2k^2h)$ | $e^{10hk}(hkl^2-k^2-m^2)+e^{6hk}(2kl^2h+4h^2k^2(l^2+8m^2)+2(k^2+m^2))+e^{2hk}(-3l^2kh-4h^2k^2l^2-k^2-m^2)$ | $(-kl^2)+e^{4hk}(2kl^2-4k^2l^2h)+e^{8hk}(-kl^2+4l^2k^2h)$ | $(hkl^2+k^2+m^2)+e^{4hk}(2kl^2h-4h^2k^2(l^2+8m^2)-2(k^2+m^2))+e^{8hk}(-3hkl^2+4h^2k^2l^2+k^2+m^2)$ |
| 1,2 | $C_h ml/(k^2(e^{4hk}-1))$ | $e^{10hk}k+e^{2hk}(k+4hk^2)+e^{6hk}(-2k-4hk^2)$ | $e^{10hk}(-kh-1)+e^{2hk}(3hk+4h^2k^2-1)+e^{6hk}(-2kh+28h^2k^2+2)$ | $k+e^{8hk}(k-4hk^2)+e^{4hk}(-2k+4hk^2)$ | $(1-hk)+e^{8hk}(3hk-4h^2k^2+1)+e^{4hk}(-2hk-28h^2k^2-2)$ |
| 1,3 | $-C_h l$ | $e^{6hk}(-1)+e^{2hk}(1-4hk)$ | $e^{6hk}h+e^{2hk}(-h-4h^2k)$ | $(-1)+e^{4hk}(1+4kh)$ | $(h)+e^{4hk}(-h+4h^2k)$ |
| 3,1 | $-C_h l$ | $e^{6hk}(-1)+e^{2hk}(1+4hk)$ | $e^{6hk}h+e^{2hk}(-h+4h^2k)$ | $(-1)+e^{4hk}(1-4kh)$ | $(h)+e^{4hk}(-h-4h^2k)$ |
| 3,2 | $-C_h m$ | $e^{6hk}(-1)+e^{2hk}(1+4hk)$ | $e^{6hk}h+e^{2hk}(-h+4h^2k)$ | $(-1)+e^{4hk}(1-4kh)$ | $(h)+e^{4hk}(-h-4h^2k)$ |
| 3,3 | $-C_h$ | $e^{6hk}(k)+e^{2hk}(-k+4hk^2)$ | $e^{6hk}(-1-kh)+e^{2hk}(-3hk+4h^2k^2+1)$ | $(-k)+e^{4hk}(k+4k^2h)$ | $(hk-1)+e^{4hk}(3hk+4h^2k^2+1)$ |

## Supplementary Movies

**Supplementary Movie 1**: Dynamics of the non-motile strain (mNeonGreen, volume fraction *ϕ*_*NM*_ = 5.3 %) in mixture with a motile WT strain (volume fraction *ϕ*_*M*_ = 0.18 %). Scale bar: 50 *µ*m. The movie is 60 times accelerated.

**Supplementary Movie 2**: Time evolution of the formation of the density fluctuation in a binary mixture of a motile WT strain (mCherry, volume fraction *ϕ*_*M*_ = 0.53 %) and a non-motile strain (mNeonGreen, volume fraction *ϕ*_*NM*_ = 1.8 %). Scale bar: 50 *µ*m. The movie is 200 times accelerated.

**Supplementary Movie 3**: Dynamics of the non-motile strain (mNeonGreen, volume fraction *ϕ*_*NM*_ = 5.3 %) in mixture with a motile Δ*cheY* strain (volume fraction *ϕ*_*M*_ = 0.18 %). Scale bar: 50 *µ*m. The movie is 60 times accelerated.

## Supplementary note: Extended simulation methods

### Brownian dynamics simulations

Our simulations use a truncated and adapted version of the equation used for Brownian dynamics simulations to compute the time evolution of the positions of the mixture of motile and motile cells. The full Brownian dynamics of a set of N particles in a fluid otherwise at rest is given by :

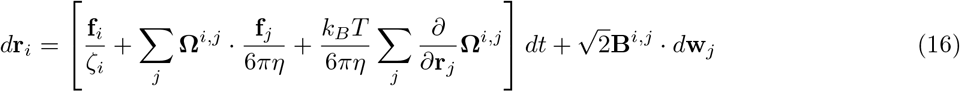

with *dt* the time step, **r**_*i*_ the positions of the particles indexed by *i*, **f**_*i*_ the forces on particle *i*, that it will also exert o<u>n</u> the fluid, *ζ*_*i*_ the Stokes drag coefficient, *η* the fluid viscosity, **Ω**^*i,j*^ the hydrodyna<u>m</u>ic-interaction tensor, and 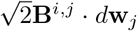 · *d***w**_*j*_ the Brownian term including random local perturbations 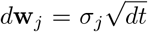, with *σ*_*i*_ a Gaussian random variable with zero mean and unit variance. The tensor **B**^*i,j*^ is coupled to the hydrodynamic interactions via the fluctuation dissipation theorem *k*_*B*_*T*(**I***/ζ*_*i*_ + **Ω**^*i,j*^*/*6*πη*)= **B**^*i,j*^ · **B**^*i,j*^

## Approximations

The full Brownian dynamics simulations of the binary mixtures would imply solving this set of equations for the motile and non-motile particles together. In our simple scheme, we only solve for the positions of the non-motile particles, the positions of the motile ones being an ansatz as explained in the Methods section. Furthermore, we simplify the treatment of hydrodynamic interactions.

The hydrodynamic-interaction tensor **Ω**^*i,j*^ accounts for hydrodynamic interactions between particles, via its non-diagonal terms **Ω**^*i,j*^ with *i, j* two particle indices, as well as for direct interactions with the boundaries of the liquid domain (in principle relevant in our case) through its diagonal terms **Ω**^*i,i*^. We neglect the latter diagonal terms, **Ω**^*i,i*^ = 0, considering that the values of the diffusion coefficient of non-motile cells and the swimming speeds of the motile ones were measured within the 40 − 50*µ*m high channel, and thus already account for the mean increase in friction due to confinement. We further make a two-body approximation for the cross-particle interaction, i.e. **Ω**^*i,≠ i*^ depends only on the positions of *i* and *j*. In this case, the divergence of the interaction tensor vanishes, 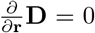. We also neglect the hydrodynamic interactions between non-motile particles, considering only the terms **Ω**^*i,j*^ **f**_*j*_*/*6*πη* for which *i* is non-motile and *j* is the flagellum or cell body of a motile cell, modeled as a force dipole. As explained in Methods, we also consider only the average flow created by the dipoles when they perform one circular revolution. We fur<u>ther n</u>eglect the effects of hydrodynamic interactions in the Brownian term, which thus reduces to 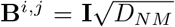, with *D*_*NM*_ = *k*_*B*_*T/ζ* the diffusion coefficient of the non-motile cells and **I** the identity matrix. Finally, the only bulk force on the non-motile cells is buoyancy-corrected gravity, since we neglect steric interactions between non-motile cells.

The simplified equation of motion for the non-motile cell *i* therefore reduces down to:

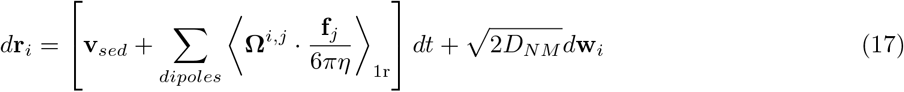

with **v**_*sed*_ = Δ*m***g***/ζ* the sedimentation speed, and the indices *j* run on the flagella and cell bodies of motile cells.

## Hydrodynamic interactions

We focus in the following on the interparticle hydrodynamic interaction term of the equation of motion (17):

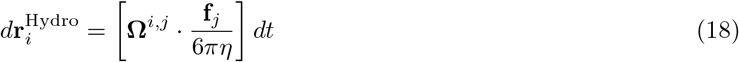

Although the interparticle hydrodynamic interaction tensor **Ω**^*i,j*^ is simply given by the Oseen tensor in unbound space, 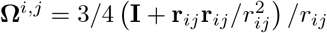, its computation is significantly more difficult and expensive in the case of confined geometries. For this, we follow an algorithm first introduced by Mucha *et al*. ^71^ and further developed by Hernandez-Ortiz *et al*. ^72^ for our geometry with periodic boundary conditions in x and y and no-slip conditions on the top and bottom walls, which allows computation in *N* log *N* time scaling. We briefly remind here the principle of the algorithm and explain the details of our implementation.

In the following, we write **f**_*j*_*/*6*πη* as “**f**_*j*_” to simplify the expressions. By definition, the flow created on particle *i* situated in **x**^*i*^ by the point force **f** ^*j*^ in **x**^*j*^ is **u**^*i←j*^ = **Ω**(**x**^*i*^, **x**^*j*^)**f** ^*j*^. Expliciting the vector and tensor components, with *α, β* = 1, 2, 3 the indices for the dimensions of space, we have:

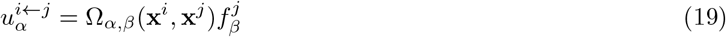

with an implicit sum over spatial indices. In our geometry, the interaction tensor can be written as a Fourier series:

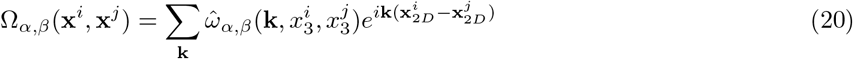

with **k** = (*l, m*) the wave vector and **x**_2*D*_ = (*x*_1_, *x*_2_) the position in the horizontal plane. The vertical position *x*_3_ is confined between the two walls in 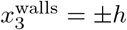, so that the channel height is *H* = 2*h*. The Fourier-transformed interaction tensor 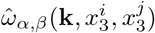 is the Green function that is solution of the partially Fourier-transformed Stokes equation for a point source:

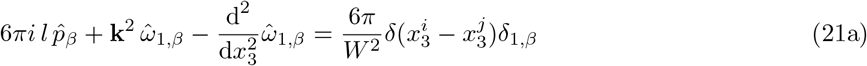

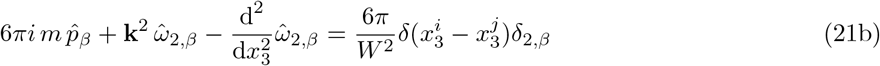

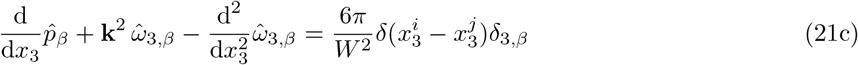

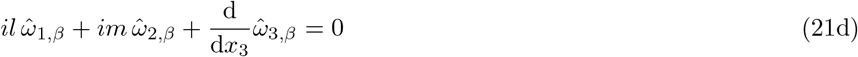

Here, 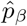 is the normalized contribution to the pressure field of the force component in direction *β, W* ^2^ is the horizontal area of the simulation box, the derivatives are taken relative to 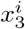, and Eq. (21d) is the incompressible flow constraint.

The gist of the contribution of Mucha *et al*. ^71^ is to realize that there is an analytical solution to this equation, which can be factorized in the following form:

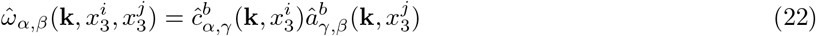

where the Boolean index *b* indicates whether the point where the flow is calculated lies below the source of flow, i.e. 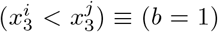. The tensors â and ĉ indeed have different analytical expressions in each case, which converge to the same well-defined value when 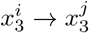. The analytical expressions of the tensors are given in a separate paragraph below. The contribution of the point force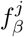 in *j* to the flow in *i* can thus be written as:

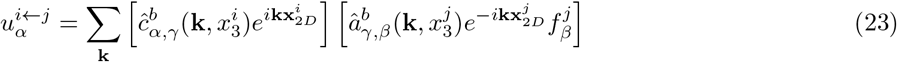

and the total flow produced by the other particles on *i* can be written as a product of contributions that respectively depend only on the source and the target:

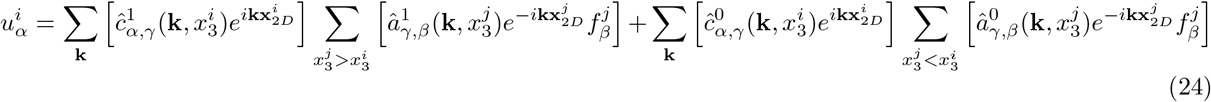

We now describe the algorithm allowing the fast calculation of these sums for all particles ^72^. At each time step, all the necessary coefficients of â and ĉ are computed. The particles are sorted according to their vertical position. The algorithm initializes a vector 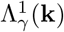, which will contain all the contribution of the cells above the current one,

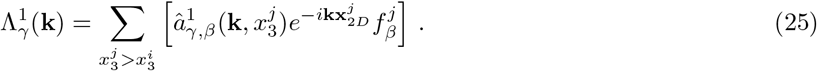

The particles are scanned from top to bottom. For each particle *i*, the contribution to the flow in **x**^*i*^ from the particles above it is computed as:

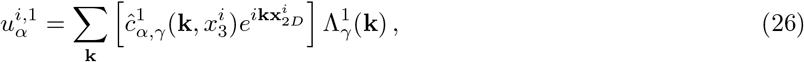

the 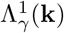 (**k**) is then updated with the contribution of particle *i*:

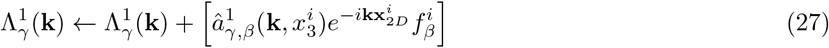

and the next particle is considered. A first top-to-bottom swipe through the particles thus allows to compute the *b* = 1 contribution to the flow for each particle. A bottom-to-top swipe similarly calculates the *b* = 0 contribution. This allows to compute the hydrodynamic interactions in *n*_*k*_*N* time, with *n*_*k*_ the number of wave vectors contributing to the sum over **k**. Importantly, this sum can be truncated because the large |**k**| have vanishing contributions to the sum. The sorting of the cells then imposes a global scaling of the computation time as *N* (*n*_*k*_ +log *N*), which is much faster than the typical *O*(*N* ^2^) scaling for brute force computing hydrodynamic interactions.

In our implementation of the algorithm, the two following modifications to the scheme are made. First, the force 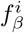 is replaced by a Gaussian-smoothed force 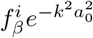, with a_0_ a cutt-off length. This is equivalent to replacing the point force by a Gaussian distribution of width *a*_0_ in the horizontal plane. It accelerates the convergence of the sum of the Fourier components by regularizing the slowly converging contributions of some of the terms for which *l* = 0 or *m* = 0, with **k** = (*l, m*), and regularizes the divergences at **x**_*i*_ = **x**_*j*_ without affecting the far field solution. This also allows us to consider a further reduced number of components *n*_*k*_. Second, since the motile cells are located at positions 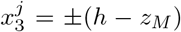, we first compute 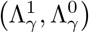 for the three cases 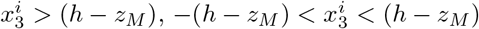 and 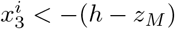 before scanning once through the non-motile particles, without the now useless sorting, to compute the total flow field they experience. Finally, dimensional analysis tells us that all distances can be expressed in units of *W* and that the hydrodynamic interaction tensor **Ω**(**x**^*i*^, **x**^*j*^) has units of an inverse distance. To avoid numerical errors due to exponential factors in the expressions of â coefficients, we define a hydrodynamic pixel size px_*H*_, such that *W* = 128 px_*H*_, and express all distances in this unit for the purpose of calculating hydrodynamic interactions, **Ω** being converted back in natural units for solving Eq. 17.

## Interaction-tensor coefficients

We remind here the mathematical expressions for the tensors 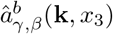 and 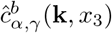 that allows computing the hydrodynamic interactions. The analytic expressions of all the coefficients were originally given in ^72^. We start with the point force contribution coefficient 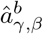. Taking into account the symmetries of the problem, for the non-zero wave vectors **k** = (*l, m*)≠ **0**, it reduces to:

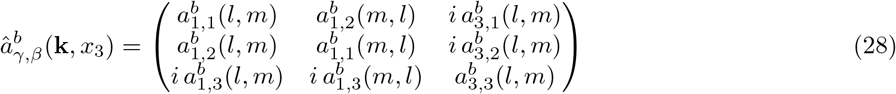

The six above-defined 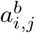 depend on *b* = (0, 1) as well as on the position *x*_3_. For wave vectors **k**≠ **0**, the later dependence takes the functional form:

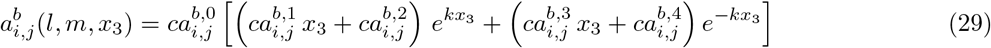

The coefficients 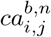 only depend on **k** = (*l, m*) and *b*. They can be calculated once at the beginning of the simulation for speed. Because several typing errors were introduced in ^72^, their correct expressions are given in Table S3.

The tensor of coefficients accounting for the position where the flow is calculated, 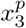, are given in the case of non-zero wave vectors **k** = (*l, m*)≠ **0** by:

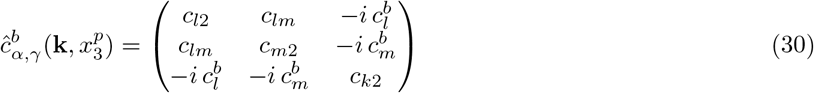

with the six distinct coefficients assuming the following values:

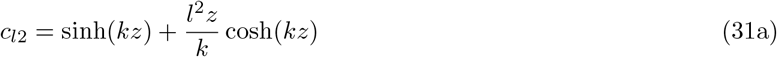

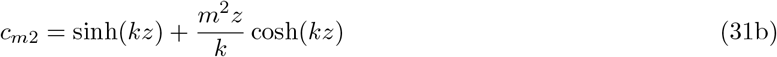

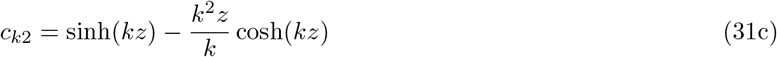

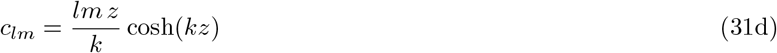

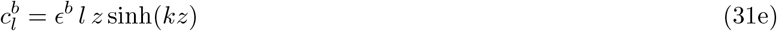

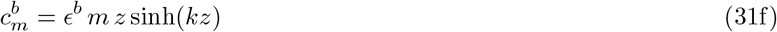

with *ϵ*^*b*^ = 1 if *b* = 1 and *ϵ*^*b*^ = −1 if *b* = 0, and the variable 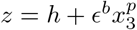.

In the case of the contribution of the **k** = **0** terms to the flow, which express the no-net-global-flow condition, the two tensors are given by:

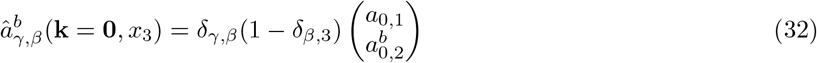

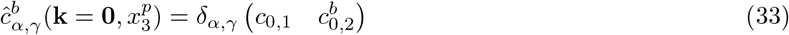

with 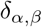 being the Kronecker delta. The two vectors form a scalar product when 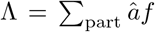and ĉ are multiplied to compute the flow in Equation (26), such that the **k** = **0** contribution to the flow for a pair-wise interaction is

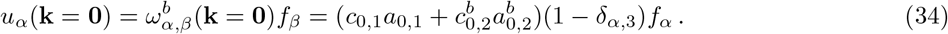

Note that the contribution of the vertical component *f*_3_ is zero for **k** = **0**, reflecting the no-flow boundary condition at the walls. Finally, the coefficients *a*_0,*n*_ and *c*_0,*n*_ are given by:

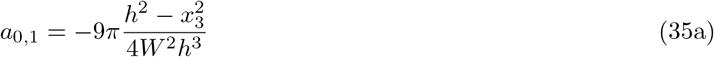

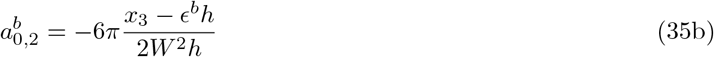

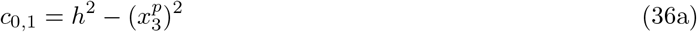

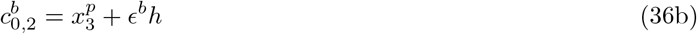

with *ϵ*^*b*^ = 1 if *b* = 1, *ϵ*^*b*^ = −1 if *b* = 0, *h* the half-height and *W* ^2^ the horizontal area of the simulation box, as defined previously.

